# Characterizing Adeno-Associated Virus Capsids with both Denaturing and Intact Analysis Methods

**DOI:** 10.1101/2023.06.20.543103

**Authors:** Jack P. Ryan, Marius M. Kostelic, Chih-Chieh Hsieh, Joshua B. Powers, Craig A. Aspinwall, James N. Dodds, John E. Schiel, Michael T. Marty, Erin S. Baker

## Abstract

Adeno-associated virus (AAV) capsids are among the leading gene delivery platforms used to treat a vast array of human diseases and conditions. AAVs exist in a variety of serotypes due to differences in viral protein (VP) sequences, with distinct serotypes targeting specific cells and tissues. As the utility of AAVs in gene therapy increases, ensuring their specific composition is imperative for correct targeting and gene delivery. From a quality control perspective, current analytical tools are limited in their selectivity for viral protein (VP) subunits due to their sequence similiaries, instrumental difficulties in assessing the large molecular weights of intact capsids, and the uncertainity in distinguishing empty and filled capsids. To address these challenges, we combine two distinct analytical workflows that assess the intact capsids and VP subunits separately. First, charge detection-mass spectrometry (CD-MS) was applied for characterization of the intact capsids and then liquid chromatography, ion mobility spectrometry, and mass spectrometry (LC-IMS-MS) separations were used for capsid denaturing measurements. This multi-method combination was applied to 3 AAV serotypes (AAV2, AAV6, and AAV8) to evaluate their intact empty and filled capsid ratios and then examine the distinct VP sequences and modifications present.

---

Gene therapy has recently emerged as a promising treatment for human diseases and conditions because it is able to alter a patient’s genetic code and address the root cause of many disorders.^1, 2^ Current gene therapy approaches use a variety of biological vectors including bacteria, viruses, and plasmid DNA to deliver the genetic payload that can either replace the faulty gene, add genes, or silence the mutated gene. However, because of these alterations, gene therapy is approached with great caution and requires extensive analytical characterization and validation of these vectors to ensure patient safety and effective treatment.

One viral gene therapy vector gaining significant traction is the adeno-associated virus (AAV). AAVs are favored for gene therapy due to their relatively low structural complexity and relatively high immune system tolerability^4^. Structurally, AAVs are icosahedral higher order protein structures as shown in **Figure 1A**. Each AAV is comprised of three different viral protein (VP) subunits, VP1 (≈81 kDa), VP2 (≈65 kDa), and VP3 (≈60 kDa), configured in roughly a 1:1:10 ratio (VP1:VP2:VP3).^2,5^ An intact AAV capsid is thus on the megadalton (MDa) scale, capable of housing an ≈5 kb DNA genome. AAV capsids can deliver and insert genetic payloads into a variety of cell types that are contingent on the specific AAV serotype, which in turn differ by the VP sequence subunits. For example, a common AAV serotype is, AAV2 that is used to target retinal dystrophy (tradename Luxturna) and AAV9 for the treatment of spinal muscular atrophy (tradename Zolgensma).

**Figure 1.**
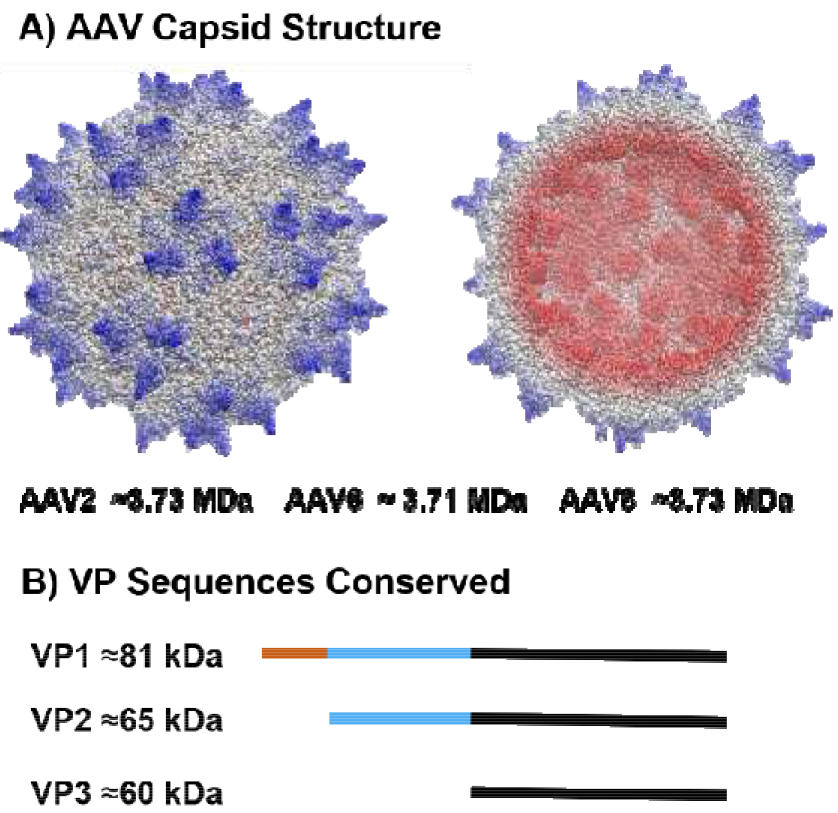
AAV capsid and sequence information. **A)** Intact AAV capsid structure, showing exterior and interior. Intact AAV capsids have an icosahedral structure formed from three subunit viral proteins (VP1, VP2, and VP3). Experimentally observed masses of empty capsids for AAV2, AAV6, and AAV8 are shown below the capsid structure. **B)** VP sequences are conserved within a given AAV serotype as shown using colored bars. The entire VP3 sequence (black) is contained within the sequence of VP2, with the additional sequence portion shown in blue. The VP2 sequence (blue and black) is also contained within the sequence of VP1, with the unique N-terminus shown in orange. Approximate masses of each VP regardless of serotype are also noted.

The structural composition of each AAV, including its constitutive subunits, genetic payload, and their appropriate combination, determines its biological function. Assessing the quality and characteristics of AAV is vital, yet this assessment presents several analytical challenges. First, the VP subunits are susceptible to post-translational modifications (PTMs), such as phosphorylation, deamidation, and oxidation, which affect the AAV’s stability, tissue tropism, and transduction efficiency.^6^ Second, there is a high degree of sequence overlap between VP1, VP2, and VP3 because the sequence of VP3 is contained within the sequence of VP2 and because VP2 is contained within VP1 (**Figure 1B**). Thus, VP1 and VP2 have unique N-terminal sequence region but share common C-terminal sequences, which limits bottom-up proteomic analysis.^5, 6^ Furthermore, the large MDa mass of each intact AAV capsid presents a unique analytical barrier for intact characterization using traditional native mass spectrometry. This factor is further complicated by the DNA genome, thus necessitating methods for discerning the degree of encapsidation.

Currently, AAV serotype identification and characterization methods use a variety of analytical techniques, including enzyme-linked immunosorbent assay (ELISA), ultracentrifugation, electron microscopy (EM), and for the native virus, polymerase chain reaction (PCR).^9-13^ Although each method is successful in its respective aims, each has shortcomings and limitations. AAV serotype identification by ELISA is limited by a lack of serotype-specific antibodies, as well as limited specificity in distinguishing between serotypes that possess a high degree of sequence similarity.^10^ Additionally, although PCR requires smaller sample volume than ultracentrifugation (≈20 μL final volume), it is primarily used to assess genome titer rather than performing definitive sample characterization.^10,11^ In addition to the challenges of serotype identification and analysis, encapsidation presents its own unique issues. Ultracentrifugation is capable of distinguishing between empty and filled AAV capsids, but it requires large sample volumes, typically 400 μL or greater.^10^ Although EM requires less sample volume than ultracentrifugation, EM traditionally requires highly purified samples for accurate component characterization. Furthermore, as EM is an image-based technique, it requires visual assessment of each image from a trained technician, limiting its utility in high-throughput applications.

Recently, mass spectrometry (MS) has been used to characterize both intact AAV capsids and their denatured VP subunits across a variety of serotypes. For intact AAV capsid analysis, charge detection (CD)-MS has alleviated several analytical challenges faced by traditional MS-based approaches.^5, 14-17^ CD-MS is a single-particle MS technique where the masses of individual ions are obtained by the simultaneous measurement of their mass-to-charge (*m*/*z*) ratios and charge.^18-20, 21^ This approach provides accurate mass determinations of high molecular weight samples. Here, we use to analyze intact empty and filled AAV capsids, which are on the MDa scale. This measurement is challenging with traditional MS methods due to the overlapping signals between charge states of large molecules, the inability to distinguish isotopic distribution within an isolated charge state, and instrumental limitations in the *m*/*z* an detector ranges.^22-25^ CD-MS was first demonstrated on home-built instruments^14^ and has been recently implemented on commercial Fourier Transform (FT) platforms (e.g. Orbitraps).^21, 26-28^ For example, Ebberink *et al*. were able to distinguish between empty and filled AAV capsids based on their genetic payload content using an ultra-high mass range (UHMR) Orbitrap mass analyzer.^29^

Although early Orbitrap CD-MS methods provided promising results, initial datasets were limited by high charge uncertainty, ions that did not survive the whole transient, and suffered from low signal-to-noise ratios. To address these limitations, a recent study by Kafader et al. highlighted a new methodology termed selective temporal overview of resonant ions (STORI) CD-MS. STORI CD-MS has benefits over other approaches, including the correction of intermittent signals as well as the differentiation of single and multiple ions at the same frequency. STORI CD-MS therefore enables improved peak shape and resolution compared to conventional CD-MS^5, 17, 30, 31^, as well as the capability to detect partial and overfilled capsids. However, recently, the Heck lab has developed a frequency tracking method for conventional CD-MS, which has been shown to improve resolution for various analytes, but we did not have access to the transient data for this manuscript. ^21^

Denatured VP subunit analysis is also of great importance. A previous method by Liu *et. al*. was used to denature and rupture the intact AAV capsid and then chromatographically separate individual VP subunits for detection with fluorescence and MS.^6, 32^ This subunit analysis method provided the unique selectivity for VPs of a variety of AAV serotypes. Additionally, VPs with post-translational modifications (PTMs), including oxidation and phosphorylation, were also distinguished using this approach.

Here, we build on these methods by adding drift tube ion mobility spectrometry (DTIMS) with LC-MS measurements for top-down structural characterization of the AAV VP subunits and their corresponding PTMs. DTIMS provides gas-phase separation of molecules based on their size, shape, and charge,^33, 34^ which enables a complementary assessment of the gas phase size of VP subunits both with and without PTMs. We then used both an LC-DTIMS-MS evaluation of individual VP subunits as well as the STORI CD-MS evaluation of the intact capsids to help provide a more comprehensive analytical evaluation of AAVs on the intact and VP subunit levels. These combined methods can address many of the remaining analytical challenges associated with AAV characterization and offer the ability to improve quality control of AAVs from intact formulations to the VP subunit level.

## Experimental

### Materials and Methods

AAV formulations were purchased from Virovek (Hayward, CA). AAV formulations included empty and filled capsids of serotypes 2, 6, and 8, where filled capsids contained the cytomegalovirus (CMV) promoter region tagged with green fluorescent protein (GFP) functioning as the reporter, corresponding to an expected mass of 0.783 MDa.^35^ AAV samples were also purchased from Virovek, wherein each sample was shipped and stored in phosphate buffered saline (PBS) with Pluronic F-68 at a volume of 0.001%. AAV2 capsid storage solutions also contained sodium citrate buffer to prevent aggregation at high concentrations. Solvents and mobile phase additives were purchased as LC-MS grade reagents from Millipore Sigma (Burlington, MA).

### Intact AAV Preparation

All intact AAV capsids were buffer exchanged into 0.2 mol/L ammonium acetate and were concentrated using a 100 kDa molecular weight cut-off filter. Each sample was buffer exchanged at least twice consecutively to ensure a total replacement of the AAV sample from the PBS based storage solution to the ammonium acetate to minimize contamination of the source region.

### Conventional Orbitrap CD-MS

Conventional Orbitrap CD-MS, pioneered by the Heck lab and Alexander Makarov^36, 37^, was performed on an UHMR Orbitrap (Bremen, Germany) as described previously. ^28, 29, 31, 37^ Conventional Orbitrap CD-MS uses the signal intensity or the signal-to-noise (S/N) ratio of single ions to determine their charge.^31^ For our measurements, we used the S/N ratio to determine the charge of single ions. The S/N to charge slope conversion was calibrated using standard proteins with known charge states as previously described.^31^ Similar to our previous work, we found a slope of 0.205 S/N per charge at 240k resolution and a slope of 0.148 S/N per charge at 120k resolution worked well for these experiments. Instrument parameters were the same between conventional CD-MS and STORI CD-MS, except the enhanced Fourier transform (eFT) was left on and the data was centroided and the noise threshold was set to 0.

### STORI CD-MS

STORI CD-MS was performed on an UHMR Orbitrap (Bremen, Germany) as described previously by Kafader and coworkers using a beta version of the Direct Mass Technology mode (Thermo Scientific).^30^ Briefly, STORI tracks the intensity of a single ion during its transient as a function of time. The slope of the resulting time versus intensity plot for a single ion is directly related to the charge of the ion. ^30, 36^ STORI slopes in this work were calibrated by resolving intact AAV capsids with native MS and overlaying the STORI CD-MS data over the native MS data to confirm the m/z values and their associated charge states prior to STORI CD-MS analysis. For 240k resolution, a STORI conversion of 112100 slope per charge was used, and for 120k, a STORI conversion of 113800 slope per charge was used. STORI mode was enabled with the raw file, and the m/z range was 15,000 to 40,000. Argon was used as the collisional gas, and trapping pressure was set to 10. Importantly, setting the bent DC flatapole to 10 V improved transmission of filled AAV capsids. The spray voltage was variable from 1.0–2.0 kV. For empty capsids, the resolution was set to 240k resolution, and for filled capsids it was set to 120k resolution. Lowering the resolution allowed the acquisition of more scans and thus higher ion counts for the filled AAV capsids. For empty capsids, the DC offset voltage was set to 10V to 21 V, but for filled capsids this was lowered to 0 V.

STORIboard software (Thermo Scientific and Proteinaceous) was used to convert the raw files to direct mass technology (DMT) For processing the raw STORI files, we started with a default template for analytes over 500 kDa. For the STORI processor, the r-squared threshold was set to 0.99, with a duration threshold of 3, time of death 0, maximum time of birth of 0, and signal to noise threshold of 0, and these settings should retain 100% of the signal ions for a 512 ms transient time. The charge assigner parameters were kept at the default of 50 *m/z* tolerance and number of averaged ions 50. Three replicate measurements were taken for each AAV to allow for standard deviations so that error bars can be used. Subsequently, the STORI files were then processed in UniDecCD, a free software available from https://github.com/michaelmarty/UniDec/releases.^31, 38^

### UniDecCD Processing

UniDecCD (UCD) was used to process and plot conventional Orbitrap CD-MS data and STORI CD-MS data. The *m/z* bin size was set to 400 and a charge bin size of 1 and mass bin size of 1000 Da were used. For conventional Orbitrap CD-MS, the *m/z* gaussian smoothing was set to 1. For conventional CD-MS deconvolution, the smooth nearby points was set to 1, the *m/z* full-width-half-maximum (FHWM) was set to 10 and the charge FWHM was set to 8, similar to our previous work.^31,39^ Because STORI has lower charge uncertainty compared to other Orbitrap based CD-MS methods^31^, no deconvolution was used for the STORI AAV data, but the maximum mass was set to 7 MDa.

### LC-IMS-MS Parameters

Chromatographic parameters for AAV characterization were adapted from Liu et al. to optimize on-column separation of the AAV VP subunits using an LC-IMS-MS platform.^6^ Briefly, a 1290 Agilent UPLC (Santa Clara, CA) was used with a Waters glycoprotein BEH Amide Column (2.1 mm x 150 mm, 1.7μm). 7 μL injections were performed, where mobile phase A was comprised of water with a volume fraction of 0.1% trifluororacetic acid (TFA)with 5mM ammonium acetate, and mobile phase B was a volume fraction of 0.1% TFA in acetonitrile with 5mM ammonium acetate. The LC column temperature was maintained at 60 °C with a 0.2 mL/min flow rate. For extended chromatographic conditions and gradient information, see **Table S1** in the supplemental information document. Following chromatographic separation, AAV VP subunits were analyzed using an Agilent 6560 IM-QTOF (Agilent Technologies, Santa Clara, CA), which has been previously characterized in detail. Briefly, in this study the subunits were ionized using the Jet Stream ESI source operated in positive ion mode with source conditions detailed in **Table S2** in the supplemental information document. Following ionization, analytes are pulsed into a uniform field drift tube with a uniform electric field drop of ≈17 V/cm for mobility separation prior to entering the time-of flight (TOF) mass analyzer. The extended IM-QTOF parameters are also detailed in **Table S3** in the supplemental information document.

Following acquisition, the Agilent .d data files were converted to .mzml files using MSConvert^41^, and uploaded to Universal Deconvolution software (UniDec)^38^ for analysis and visualization. A detailed description of the file conversion process of Agilent .d data files and their analysis using UniDec can be found in the supplemental file titled “UniDec Deconvolution of Agilent .d Data”. UniDec enables the deconvolution of complex mass and IMS spectra, as well as charge dimension visualization and IMS calculations such as profiling collision cross section (CCS) values. UniDec’s method for deconvolution employs an iterative Bayesian deconvolution to determine the effective contribution of each *m/z* and charge to the overall spectrum, which has been described in previous publications.^38^ The specific UniDec parameters used to process, deconvolve, and visualize the data can be found in **Table S4**. All data files are located at ftp://massive.ucsd.edu/MSV000092048/ and https://massive.ucsd.edu/ProteoSAFe/dataset.jsp?accession=MSV000091650 for LC-IMS-MS and CD-MS experiments respectively.

## Results and Discussion

### CD-MS Empty and Filled AAV Capsid Analysis

Because they are used as gene therapeutic vectors, it is critical to determine the successful encapsidation of intact AAV capsids that are filled with the desired payload. In this instance, the genetic payload was CMV-GFP, which is comprised of 2,544 bases and has an expected mass of 0.783 MDa.^35^ First, analysis of intact AAV capsids was performed on empty AAV2, AAV6, and AAV8 capsids using both conventional Orbitrap CD-MS and STORI CD-MS. For the empty capsids, AAV2 and AAV8 had similar masses of ≈3.70 MDa and AAV6 had the lowest mass (≈3.63 MDa) as shown in **Figure 2A**. With STORI CD-MS, the mass peaks for empty AAVs had full-width half-maximums (FWHMs) of 230 kDa to 300 kDa, which was similar in resolution to the analysis of AAV8 capsids also purchased from Virovek and analyzed in a previous study by Jarrold and co-workers using a custom-built high-resolution CD-MS instrument.^35^

**Figure 2.**
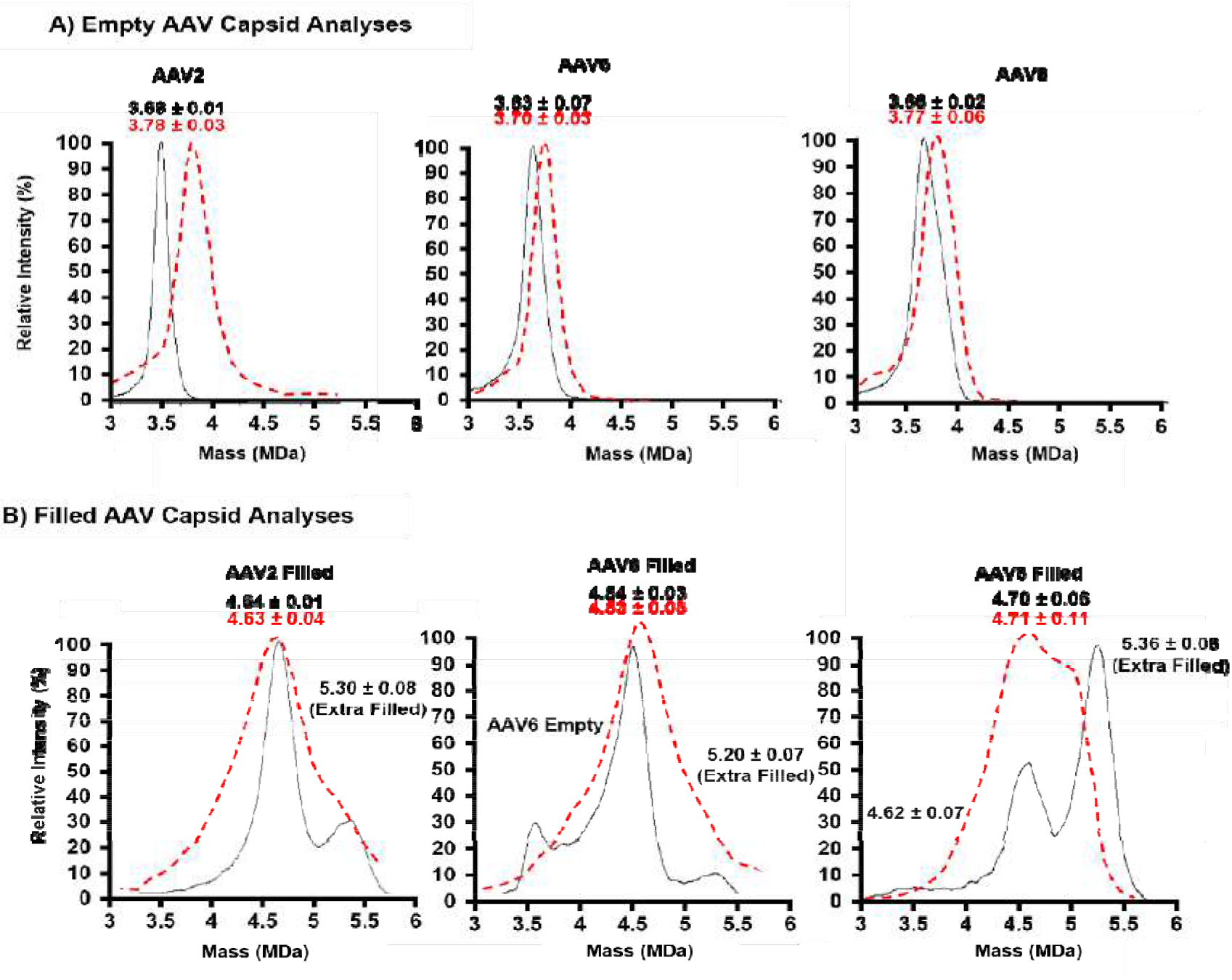
Comparing the resolving power of CD-MS and STORI CD-MS. **A**) The peaks and masses of empty AAV2 (left), AAV6 (middle), and AAV8 (right) capsids for traditional Orbitrap CD-MS (red dashed) and STORI CD-MS (black) deconvolution. STORI CD-MS narrowed peak shapes and lowered the masses when compared to CD-MS. **B)** The peaks and masses for CMV-GFP filled AAV2 (left), AAV6 (middle), and AAV8 (right) capsids were also assessed. Again, STORI CD-MS shows improved peak shape, distinguishing multiple peaks for empty and filled capsids.

Differences were observed in the deconvolved conventional Orbitrap CD-MS and STORI CD-MS mass peaks. STORI CD-MS had narrower peak widths over conventional CD-MS. The narrower peak widths were likely due to the capability of STORI CD-MS to have more accurate charge assignments with single ion by tracking the transient of each Ion rather than conventional Orbitrap CD-MS, which typically only uses the post Fourier transform data. The more precise charge assignments from STORI result in a lower charge uncertainty compared to conventional Orbitrap CD-MS, although new frequency chasing methods may further improve the resolution of conventional Orbitrap CD-MS.^17^ Interestingly, STORI gave lower masses of empty AAV capsids compared to conventional Orbitrap CDMS, but the masses matched more closely for the filled AAV capsids (Figure 2). The observed lower masses are possibly due to shifted AAV frequencies caused by collisions with background gas in the Orbitrap during the longer transient measurements, which gives an arbitrarily lower charge, and this has similarly been observed for conventional Orbitrap CDMS AAV measurements at higher transient times.^17^ The filled AAV capsid masses from both STORI and conventional Orbitrap CDMS match more closely because those were collected with a lower transient time, which lowers the resolution and thus lowers the probability that an AAV collides with background gas in the Orbitrap and shifts in frequency. It is important to note that these CD-MS measurements were collected on a high frequency (HF) UHMR and may not be observed on other platforms.

Following analysis of the empty AAV capsids, filled AAV capsids loaded with the promoter region of CMV-GFP were evaluated with conventional CD-MS and STORI CD-MS (**Figure 2B**). The filled capsids for each serotype possessed an additional 0.9 MDa mass shift compared to the empty AAV capsids, accounting for the CMV-GFP, which has a theoretical shift of 0.783 MDa. All serotypes also had an additional mass shift of 0.7 MDa, illustrating extra filling, potentially representing an additional intact second cargo, or a truncated version of the cargo.^36^ AAV2 and AAV6 capsids possessed a small contribution of the extra filled population, but a very high abundance of the extra filled distribution was noted for AAV8. For AAV8, this result may suggest that the preparation favors a capsid with two intact, or one intact and one truncated DNA cargos, which has been reported previously.^35,42, 43^

As with the empty capsids, in the conventional CD-MS to STORI CD-MS comparison, there is also a clear improvement in resolution with STORI CD-MS. For the AAV mixtures, STORI CD-MS allowed for the separation of AAV capsids containing a single payload from extra filled capsids containing more than a single payload. Additionally, for AAV6, STORI CD-MS was also able to distinguish a population of capsids corresponding to the unfilled AAV6 mass, as shown in **Figure 2B**.

For comparison, both the experimental and theoretical intact capsid masses are shown in **Table 1**. Generally, the errors for both conventional CD-MS and STORI CD-MS were relatively close, with all values for STORI CD-MS generally correlating by less than 1000 ppm error from the proposed literature value, and conventional CD-MS was also less than ≈1500 ppm except for filled AAV8, which had a much higher ppm error for the filled AAV8 formulation specifically. However, because literature is limited for the filled AAVs, their mass values were calculated using the theoretical 1:1:10 ratio of VP1:VP2:VP3, and assuming that capsids were comprised of 60 VP subunits in total. Therefore, some error could occur due to these assumptions.

**Table 1:** Experimentally determined masses and percent error for both empty and filled AAV capsids acquired using both conventional CD-MS and STORI CD-MS.

| Capsid Type | Theoretical Mass (MDa) | CD-MS Observed Mass (MDa) | CD-MS Error (ppm) | STORI CD-MS Observed Mass (MDa) | STORI CD-MS Error (ppm) |
| --- | --- | --- | --- | --- | --- |
| AAV2 Empty | 3.73 | 3.78 ± 0.03 | 1340 | 3.68 ± 0.01 | 1340 |
| AAV2 Filled | 4.51 | 4.63 ± 0.04 | 2660 | 4.64 ± 0.01 | 2880 |
| AAV2 Extra Filled* | 5.29 | ND* | N/A | 5.30 ± 0.08 | 190 |
| AAV6 Empty | 3.71 | 3.70 ± 0.03 | 270 | 3.63 ± 0.07 | 2156 |
| AAV6 Filled | 4.49 | 4.53 ± 0.08 | 890 | 4.54 ± 0.03 | 1113 |
| AAV6 Extra Filled* | 5.27 | ND* | N/A | 5.20 ± 0.07 | 570 |
| AAV8 Empty | 3.73 | 3.77 ± 0.06 | 1070 | 3.66 ± 0.03 | 1328 |
| AAV8 Filled | 4.51 | 4.71 ± 0.11 | 4430 | 4.70 ± 0.06 | 4213 |
| AAV8 Extra Filled* | 5.29 | ND* | N/A | 5.36 ± 0.06 | 1323 |

Extra Filled\* = Assumed to have two of the CMV-GFP payloads. ND\* = Not detected.

Because STORI CD-MS had higher resolution than conventional CD-MS,^5, 17, 31^ it was evaluated for separating empty and filled capsids for each serotype. To perform this study, a predefined 1:6 (v/v) ratio, post buffer exchange, of the empty to filled capsids for each serotype was mixed and analyzed (**Figure 3**). STORI CD-MS was able to successfully resolve empty, filled, and over filled capsids in the same sample with 3 min to 15 min acquisition times to accumulate enough ions. The AAV spectra differed in each scenario due to the varying amounts of overfilled capsids in each formulation, which is reflected by the differential relative intensities of the empty and filled capsids. Due to the v/v ratios being mixed post-buffer exchange, this led to slightly different rations in terms of concentration. However, for AAV6, the filled AAV sample had some empty AAV capsid present, leading to a higher amount in the mixed analysis of that serotype. Overall, STORI CD-MS offered sufficient resolution to distinguish different subpopulations of AAVs without additional processing or deconvolution.

**Figure 3.**
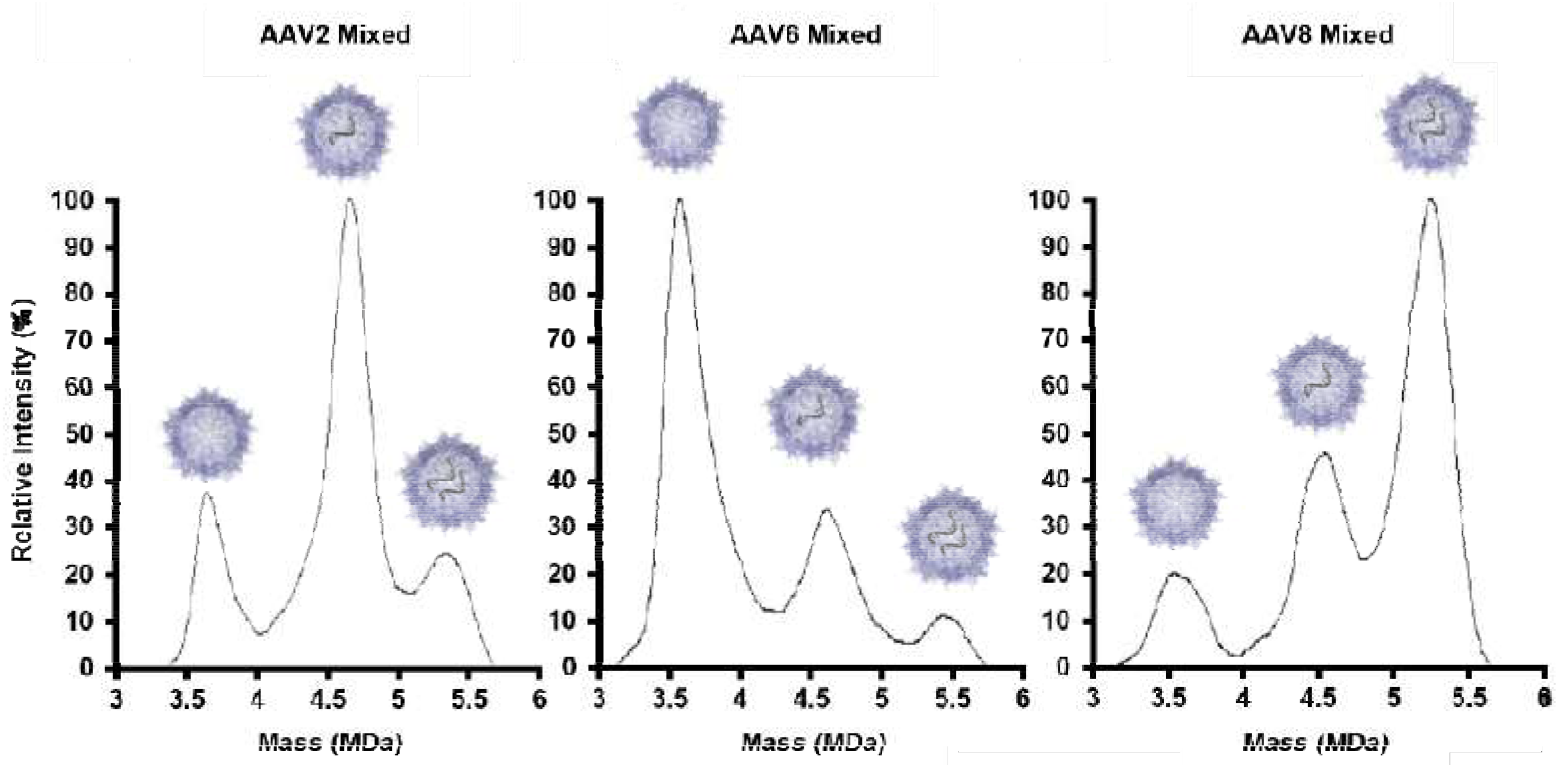
STORI CD-MS analysis of mixed empty and filled capsid preparations for AAV2 (left), AAV6 (middle), and AAV8 (right). Samples were created by mixing the empty and filled AAV capsids for each serotype at a ratio of 1:6 ratio of empty to filled, which resulted in various relative ratios of empty and filled capsids due to different over-filling amounts for each serotype.

### HILIC Separation of VP Subunits

Analysis of individual VP subunits is also a crucial step in the characterization of AAVs as gene therapeutic vectors. The high degree of overlap in the VP amino acid sequences within a given serotype necessitates advanced analytical methods capable of separating proteins with similar characteristics.^5^ The combination of LC-IMS-MS applied here is able to distinguish VP subunits within a given AAV serotype, as well as VP subunits across serotypes. The LC gradient used was the same as a previously established HILIC method,^6^ with adaptations made for the mass spectrometer to facilitate ion mobility analysis, such as the parameters outlined in **Table S2** and **Table S3** in the supplemental information document. The HILIC separations for AAV2, AAV6, and AAV8 with filled capsids are shown in **Figure 4**. Additionally, total ion chromatograms of empty AAVs are given in **Figure S1**, and an overlay of the empty and filled chromatograms shown in **Figure S2**.

**Figure 4.**
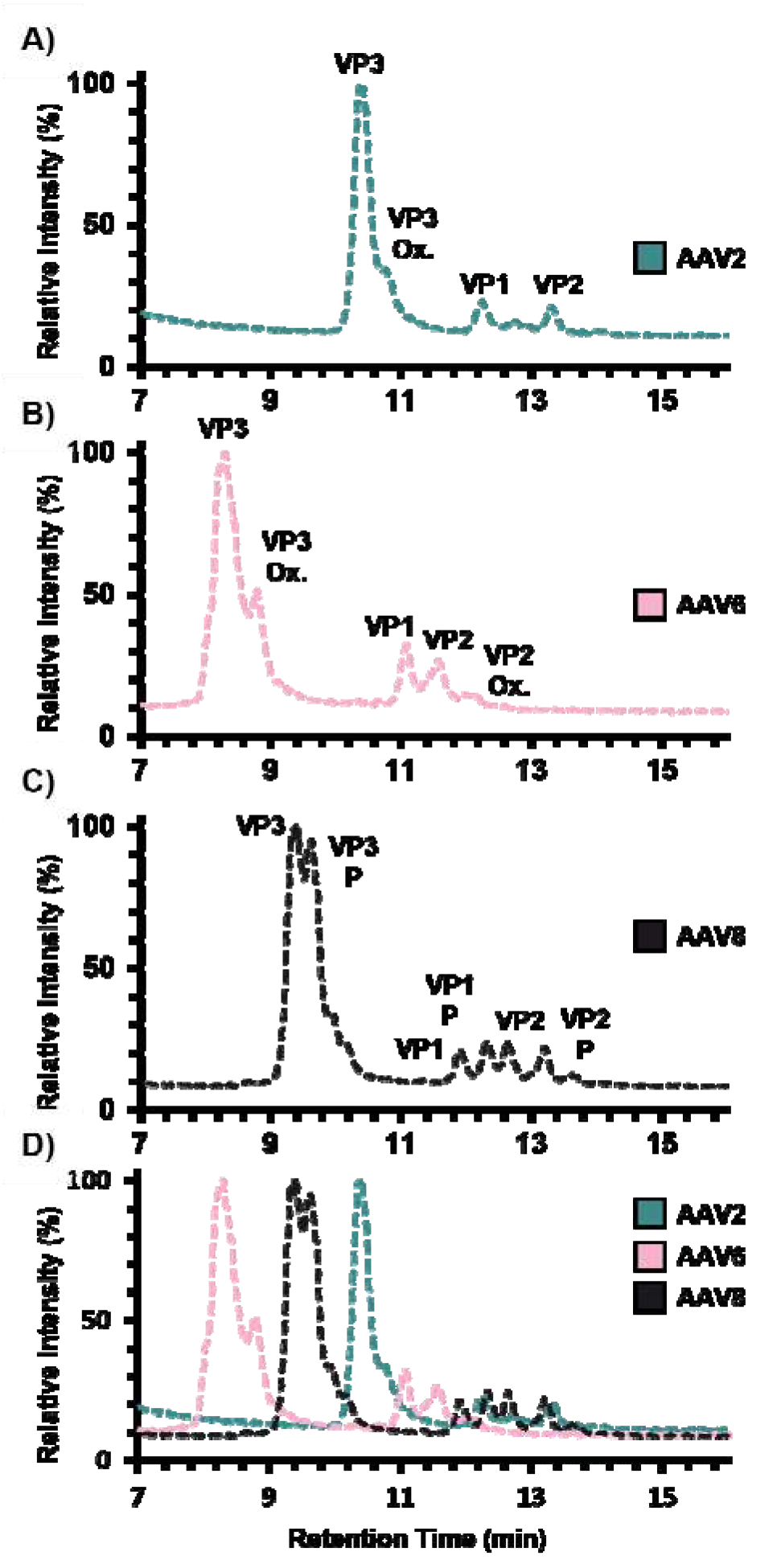
Total ion chromatograms (TICs) of denatured A) AAV2, B) AAV6, C) AAV8, and D) all three AAVs overlayed. The TICs for each AAV show separation of all three VP subunits, as well as post-translationally modified (PTM) variants. For the PTMs, oxidation of VP3 is observed for AAV2, oxidation of VP3 and VP2 occurs for AAV6, and phosphorylation of all VPs is present for AAV8. The overlay of TICs for AAV2, AAV6, and AAV8 illustrate how their VP3 have different retention times while similar LC retention times are noted for VP2 and VP3 for

For all tested AAV serotypes, the HILIC elution order was consistent with VP3 eluting first, followed by VP1, and then VP2. Consistent retention times were also observed for each VP subunits regardless of whether the sample putatively included the payload, indicating that the cargo does not impact the VP subunit method performance, though differences in intensity were observed between the empty and filled samples. The details of the bioprocess for the various samples are not known, however, it is likely that the only difference was the inclusion of transgene for the filled AAVs. Differences in overall abundance of VPs can likely be attributed to differential capsid formation in the presence of the cargo and/or slight differences in the process for the samples.

In addition to separation of VPs within a given AAV serotype, the VPs of different AAV serotypes are distinguishable through their VP3 retention times, as these provide the greatest retention time difference between the AAV serotypes in this study with elution ranging from ≈7 minutes to 11 minutes. Of the serotypes, VP3 for AAV6 eluted first, followed by AAV8, and then AAV2. VP1 and VP2, along with their PTM forms, showed very slight differences in retention time when comparing AAV6 to either AAV2 or AAV8. For AAV2 and AAV8, the retention times of VP1 and VP2 and their PTMs are very similar, with all peaks eluting in the window of ≈12 minutes to 13.5 minutes, so distinguishing between these serotypes is most easily observed by comparing VP3.

The separation of VPs with different PTMs was also observed and confirmed by MS. AAV2 and AAV6 were observed to have oxidized modifications, though other PTMs may have been present and were not detected. Specifically, VP3 was oxidized in AAV2, but both VP3 and VP2 were oxidized in AAV6. Interestingly, PTMs were observed on all 3 VP subunits of AAV8, but rather than oxidation, phosphorylation was observed. Characterization of PTMs is important because they could have implications to the overall functionality and stability of AAV formulations.^44, 45^

### DTIMS Analysis

To confirm and further characterize the PTMs and VP amino acid sequences, IMS-MS was employed following the LC separation. However, due to the complexity of the collected LC-IMS-MS spectra, advanced data analysis is also needed. Thus, it was necessary to employ UniDec^38^ to deconvolve and visualize the collected data. Prior to this study Agilent IMS-MS .d data files were incompatible with UniDec software, so UniDec was altered to accept this data. Detailed steps are included in the Supplemental UniDec Protocol. Following UniDec deconvolution, a nested drift time versus *m/z* plot is produced as shown in **Figure 5**. These nested spectra were sorted by AAV serotype across each row, and down each column by individual VPs. Interestingly, when visually comparing the nested spectra for a given VP subunit in the different serotypes, very few differences were observed. Upon confirmation with the calculated CCS values and the ΔCCS (%) calculations comparing CCS values at specific charge states for each VP in each serotype, no two CCS values were found to possess a difference greater than 0.77% (**Table S10**) in the supplemental information document. Illustrating that despite amino acid sequence differences, as confirmed by MS analysis, the overall size and structure of the VPs in the gas phase are nearly indistinguishable. Collectively noting that the ΔCCS (%) for comparisons in this work averaged ≈0.3%, separating these VP subunits by IMS alone would require ≈450 R_P_ that is currently unobtainable for most commercially available IMS platforms.^46, 47^ The high degree of similarity of the VP CCS values may potentially suggest the need for VPs to be of a similar size, regardless of sequence, to facilitate capsid formation and specific targeting. However, CCS is a gas phase descriptor of size, and future analysis of AAVs may provide more insight into VP native structure.

**Figure 5.**
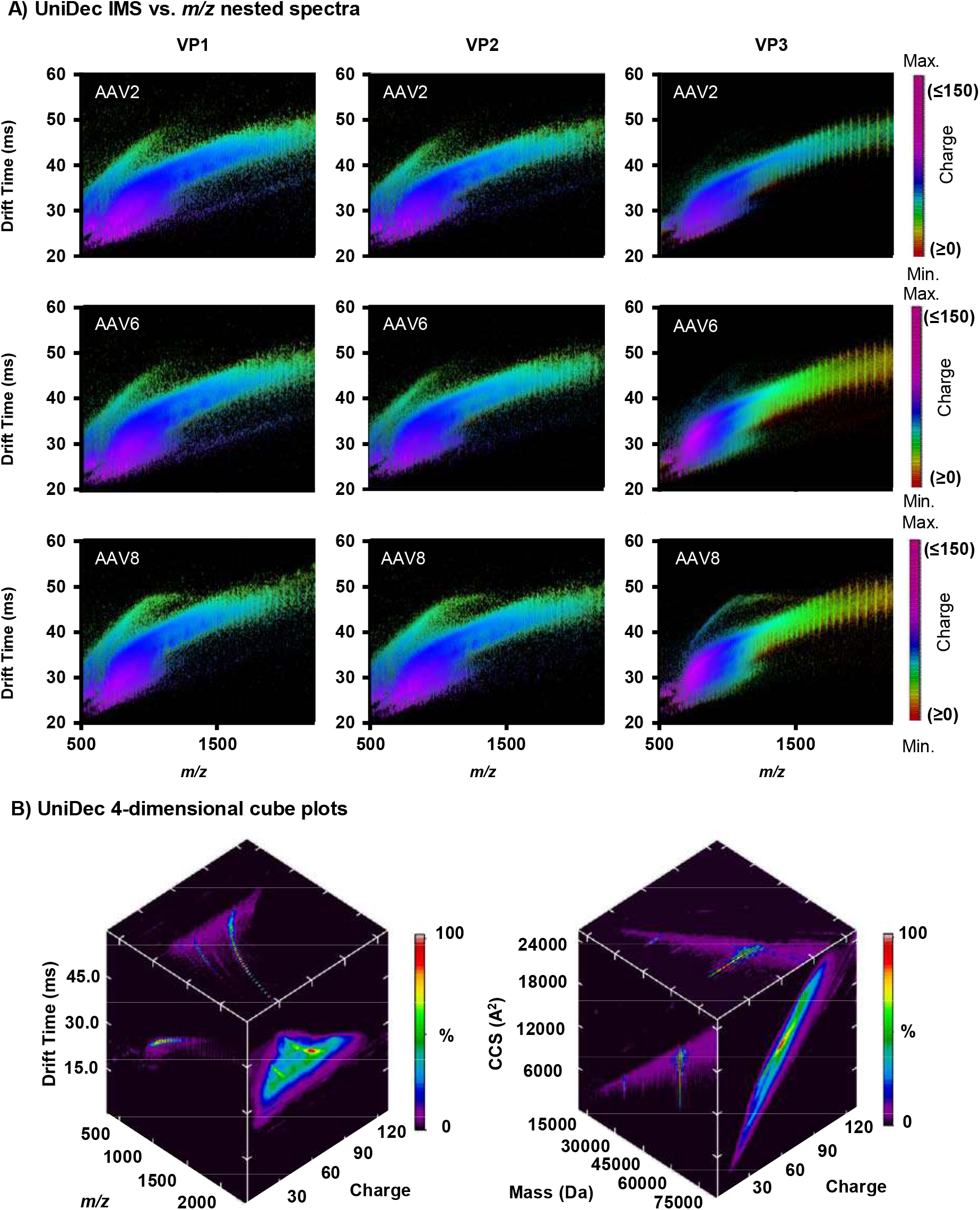
UniDec IMS-MS visualizations. **A)** UniDec IMS drift time versus *m/z* nested spectra for VP1, VP2, and VP3 across AAV2, AAV6, and AAV8. Differently colored plot areas show difference in deconvolved charge state ranging from 0 to 150+. **B)** UniDec 4-dimensional projected cube plots of AAV2 VP3 show deconvolution of raw data (left) and CCS/mass calculations (right). The UniDec cube for the raw data plots drift time (ms), *m/z*, and charge, with each face showing a different comparison, while the calculated cube plot shows CCS, mass, and charge state. Intensity for each cube is shown using a colorimetric scale with least intense represented by black and most intense shown in red.

In addition to producing charge state assigned nested spectra, UniDec also has the capability to plot and visualize data in 4 dimensions. In **Figure 5B (left)**, the cube plot shows deconvolution of raw data, with drift time on the y-axis, *m/z* on the x-axis, and charge state on the z-axis. Intensity is shown as a heat map for each comparison. Additionally, the **Figure 5B (right)** cube shows calculated values such as CCS and monoisotopic mass, allowing for rapid visualization of data in multiple dimensions. As an example of UniDec’s 4-dimensional plotting capabilities, **Figure 5B** shows cube plots for VP3 from an empty AAV2 formulation.

### MS Determination of VP and PTM Masses

Analysis of VP1, VP2, and VP3 for AAV2, AAV6, and AAV8 was facilitated through UniDec’s deconvolution and visualization of the complex spectra acquired in these experiments. Using the charge state assignment from UniDec’s deconvolution, it was possible to determine masses for each VP, as well as the modified VPs present. For both empty and filled formulations of AAV2, AAV6, and AAV8, experimentally determined VP masses agreed with the expected theoretical masses for both the modified and unmodified forms of each VP subunit.^6^ As previously noted, the type of PTM observed varied with AAV serotype. AAV2 and AAV6 had differences of ≈16 Da between modified and unmodified VP3, corresponding to an added oxygen. Although there was no VP2 modification detected for AAV2, the mass difference between modified and unmodified AAV6 VP2 was ≈32 Da, again corresponding to an oxidation PTM but with two oxygen atoms. Although oxidation was observed in both AAV2 and AAV6, for AAV8, VP1, VP2, and VP3 each has mass difference between the modified and unmodified VP masses of ≈80 Da, corresponding to phosphorylation. The observed mass differences are in agreement with previous analysis of these AAV serotypes^6^. The observed masses and mass differences for the detected modified and unmodified VP subunits across all serotypes showed no difference when comparing the empty vs. filled formulations, though it should be noted that the filled AAV capsids had a lower intensity, potentially indicating an influence of the cargo. Even with intensity differences, the similarities between filled and empty AAV capsids indicate that the cargo produced no impact on the modifications made to the subunit proteins.

### Conclusions

As the field of gene therapeutics continues to progress, it is imperative that advancements in analytical techniques used for product and process development follow suit. Here, we illustrate a multi-method workflow capable of characterizing a variety of AAV serotypes from the intact capsid level down to the VP subunits. STORI CD-MS provided accurate mass measurements of both empty and filled AAV capsids, as well as the capability to resolve multiple capsid masses in a mixture of empty and filled capsids. The varying intensities of the capsid peaks in that mixture correspond to the relative amounts of each capsid formulation, providing a metric for efficiency of product formulation, as well as the ability to discern between over or partially filled capsids. Compared to traditional CD-MS, STORI CD-MS also provided greater resolution and narrower peak shapes but gave comparable percent mass error.

HILIC chromatography paired with an IMS-MS platform achieved successful separation of VP subunits across AAV serotypes, as well as distinction of the unmodified VP peaks from any detected PTMs. The IMS dimension elucidated gas phase structural information about the VP subunits, indicating that at any given charge state, regardless of serotype, the structures of each VP were similar for the different serotypes. Mass analysis of the detected *m/z* peaks confirmed the presence of oxidation as the PTM observed for both AAV2 and AAV6, and phosphorylation for AAV8. By combining both the STORI CD-MS and LC-IMS-MS platforms for AAV analysis, it was possible to assess the quality and characteristics of the product on both an intact capsid and a VP subunit level, providing more extensive and accurate characterization of AAVs.

## Supporting information

UniDec Supplemental Tutorial

AAV Supplementary Methods

## Author Contributions

JPR and MMK both designed and carried out experimental methods, generated figures and contributed to the overall manuscript production. Conventional CD-MS and STORI CD-MS experiments and instrument maintenance were aided and facilitated by the support of CCH, CA, and MTM. LC-IMS-MS experiments were supported by JND and ESB. JES and JBP provided support in data analysis and experimental design. MTM modified UniDec code for this study. ESB, MTM, JES, and JND provided intellectual guidance and aided in the progression of these experiments.

## Disclaimer

Certain commercial equipment, software, or materials are identified in this paper to specify the experimental procedure adequately. Such identification is not intended to imply recommendation or endorsement by the National Institute of Standards and Technology, nor is it intended to imply that the equipment, software, or materials identified are necessarily the best available for the purpose.

## Acknowledgements

This work was funded, in part, by grants from the National Institutes of Standards and Technology (NIST), the National Institute for Innovation in Manufacturing Biopharmaceuticals (NIIMBL), and the National Science Foundation (CHE-1845230 to M.T.M. and CBET-2003297 to C.A.A.). Work was performed 1) within a Cooperative Agreement between NIST/ National Institute for Innovation in Manufacturing Biopharmaceuticals (NIIMBL) and 2) under a Project Award Agreement between NC State/Pfizer/NIIMBL and financial assistance award 70NANB17H002 from the U.S. Department of Commerce, National Institute of Standards and Technology.

## Notes and references

‡ Supporting information is readily available at “AAV Supplementary Methods”

§ Supplemental UniDec Protocol for analysis of Agilent IMS-MS .d data files is readily available at “UniDec Supplementary Tutorial”

§§

