## Supplementary material for "Characterizing Adeno-Associated Virus Capsids with both Denaturing and Intact Analysis Methods": UniDec Supplemental Tutorial

UniDec Deconvolution of Agilent .d Data

**Certain equipment, instruments, software, or materials are identified in this paper to specify the experimental procedure adequately.  Such identification is not intended to imply recommendation or endorsement of any product or service by NIST, nor is it intended to imply that the materials or equipment identified are necessarily the best available for the purpose.*

In this tutorial, you will learn how mass spectrometry or ion mobility spectrometry-mass spectrometry data collected on Agilent instruments can be converted and uploaded to the universal deconvolution software, UniDec, to facilitate visualization and analysis of complex spectra. In this tutorial, we will use the LC-IMS-MS data collected for the analysis of denatured AAV capsids as a case study, beginning with file conversion, and data sub-setting, ending with plotting UniDec’s 4-dimensional cube plots. Though UniDec has been coded to optimize deconvolution with minimal user intervention, it may be beneficial to get familiar with UniDec by referencing the following resources:

For a beginner tutorial, check out: <https://www.youtube.com/watch?v=e33JxgY6CJY>

Source code and additional resources can be found on the UniDec github page: [https://github.com/michaelmarty/UniDec/](https://github.com/michaelmarty/UniDec/releases)

### Getting Started with File Conversion

To start this tutorial, download the following program files at:

<https://proteowizard.sourceforge.io/download.html>

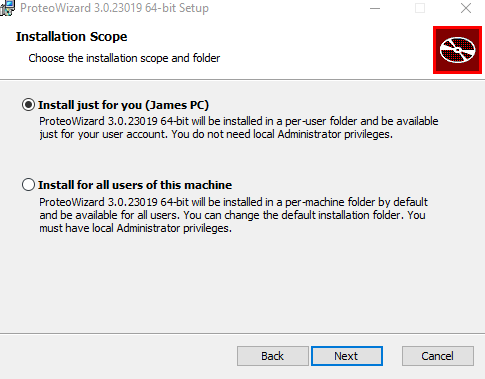
The program used to convert these files from an Agilent IMS-MS .d format to an open-source .mzML file format is MSConvert. Following the download, grant permission to install MSConvert via ProteoWizard, and choose the appropriate installation scope, for a specific user or for all users of the computer^1^.

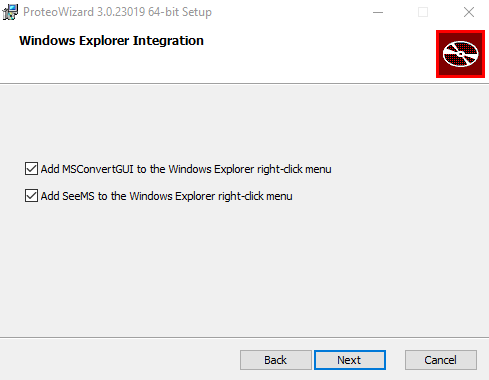
Next, choose the appropriate location destination for access to the MSConvert program as seen below:

After finalizing installation, run the program, and you should see the following window open:
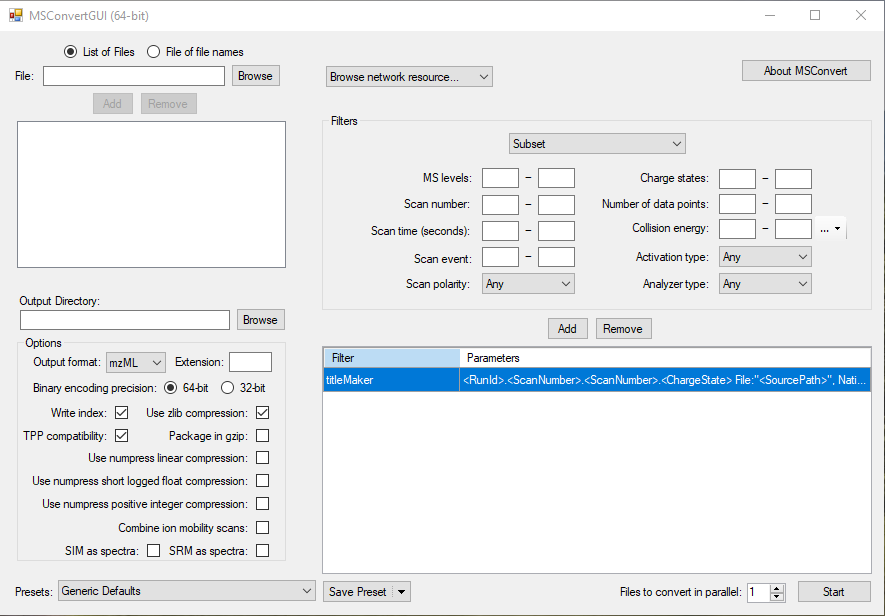

This window indicates that you have successfully installed MSConvert and the program is ready to operate. Following this step, you will need to download the demonstration data for the rest of this tutorial. To access the data that will be used for the tutorial, please download the following data file from ftp://massive.ucsd.edu/MSV000092048/: “AAV2_empty1” which itself is an Agilent .d data file. This .d file contains the data acquired during the experiments for the analysis of a denatured empty adeno associated virus capsid, in this particular instance an empty AAV2 capsid. For the remainder of this tutorial, we will focus on using UniDec to visualize our LC-IMS-MS data collected for an empty and denatured AAV2 capsid. Specifically, we will be using UniDec to facilitate the analysis of viral protein (VP) subunit 3 of AAV2.

Once you have successfully downloaded the tutorial data file, extract it by right-clicking on the folder, select “extract all”, and then proceed to open it using Agilent MassHunter’s Qualitative Analysis Navigator program (referred to as “qual” from here on out), which comes included with the purchase of an Agilent mass spectrometer. Though our data specifically contains IMS information from the experiment, we are here using qual as a means to view our data in a plot of “counts vs. acquisition time (scans).” The scan time range (in seconds) will be used to subset the data to facilitate analysis of unique peaks in the chromatogram. The normalized TIC from this data file shown below, highlights the peak we will be focusing on for analysis going forward, VP3 of AAV2.

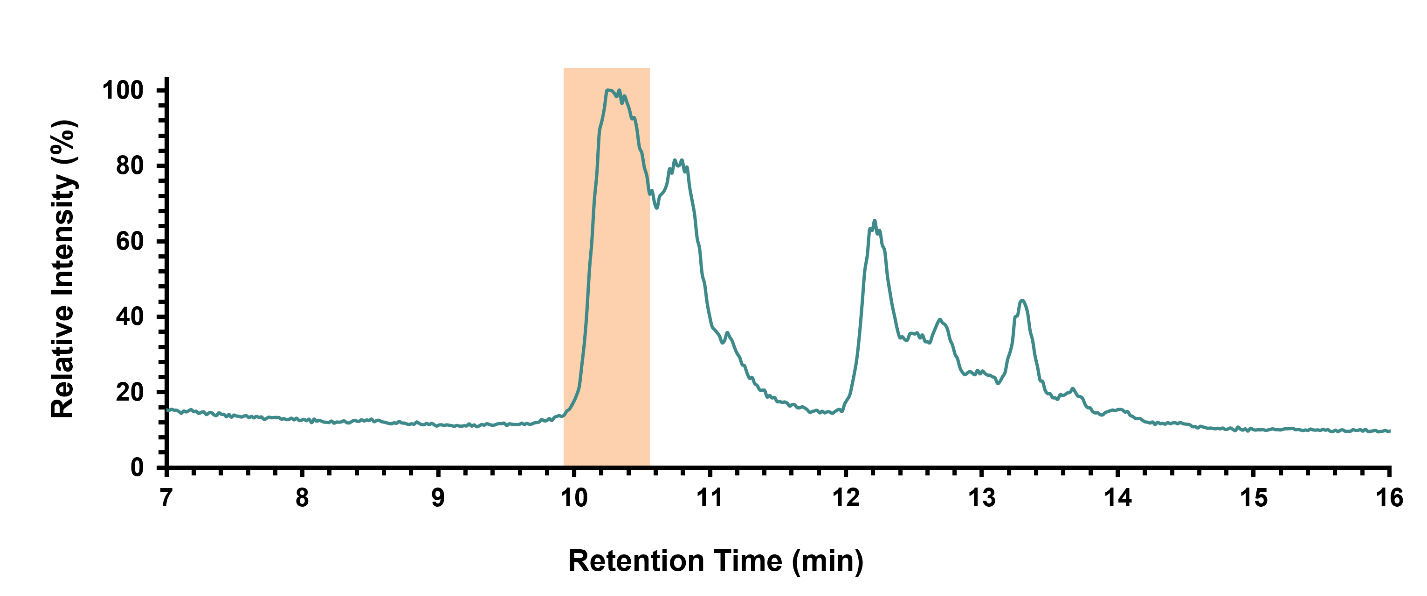

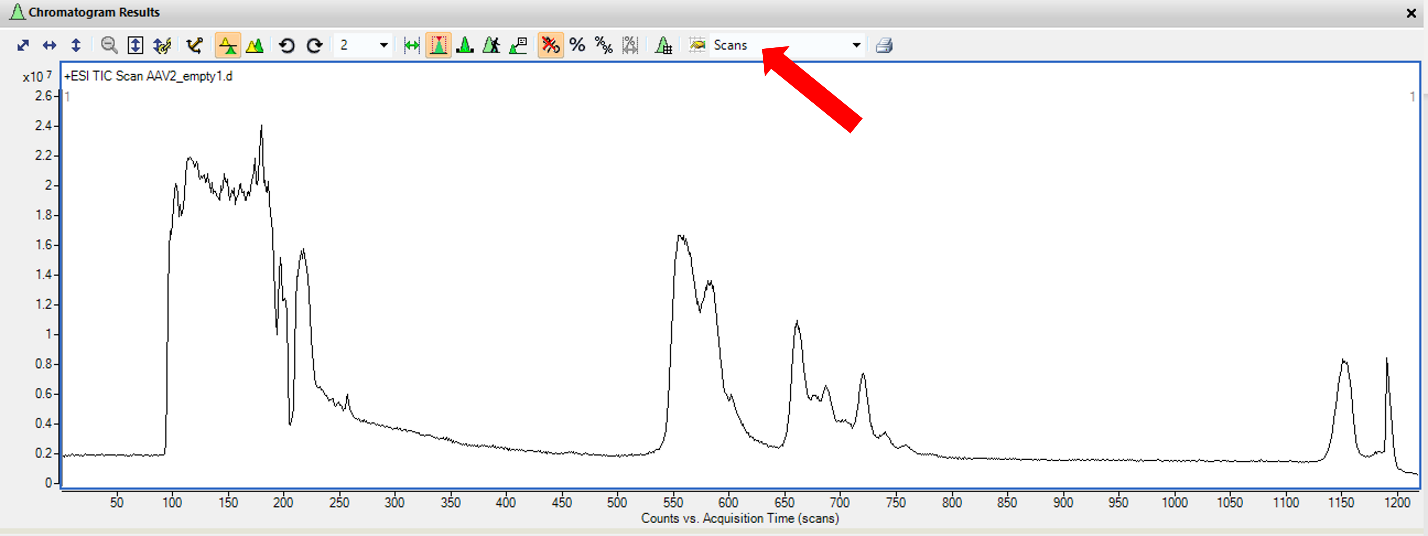
To successfully subset the data above to just include the relevant AAV2 VP3 data, switch the x-axis from “minutes” to “scans”

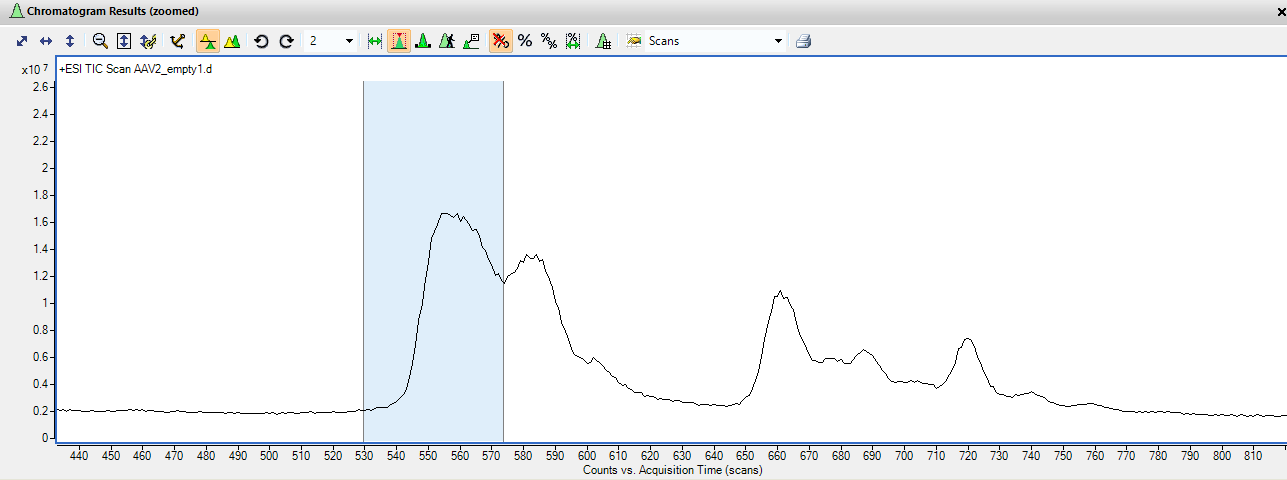
As we are focusing on AAV2 VP3, which elutes first, zoom in on that peak in qual, and record the scan times that encompass the width of the desired peak. In this instance, the peak for AAV2 VP3 has a range of ~530 to ~575 (chosen to avoid potential interference from the adjacent peak), as seen in the image below.

Now with the scan time determined, we can return to MSConvert to finally convert from a .d format to .mzML.

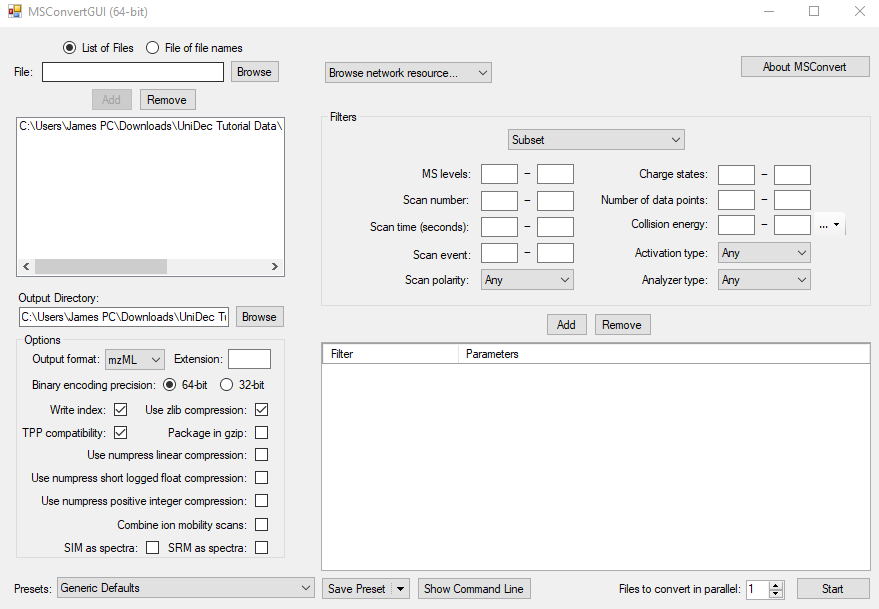
Begin by clicking the “browse” button in the top left corner of the MSConvert window, find, and select the tutorial data file, and hit the “add” button. This will upload the desired file to MSConvert for conversion. In the “output directory” name your new file in a unique and descriptive way, create the path you want your newly converted file to take. It is recommended that you create a secondary folder for the .mzML versions of your data files for organizational purposes. An important note: if you not re-name the file in the “output directory line” it will automatically be re-named the same name as the original file, even if the new .mzML file is a subset of the overall experiment. It is recommended to re-name each file (if subsetting) with the specific information about what the .mzML file contains to avoid duplicate file names.

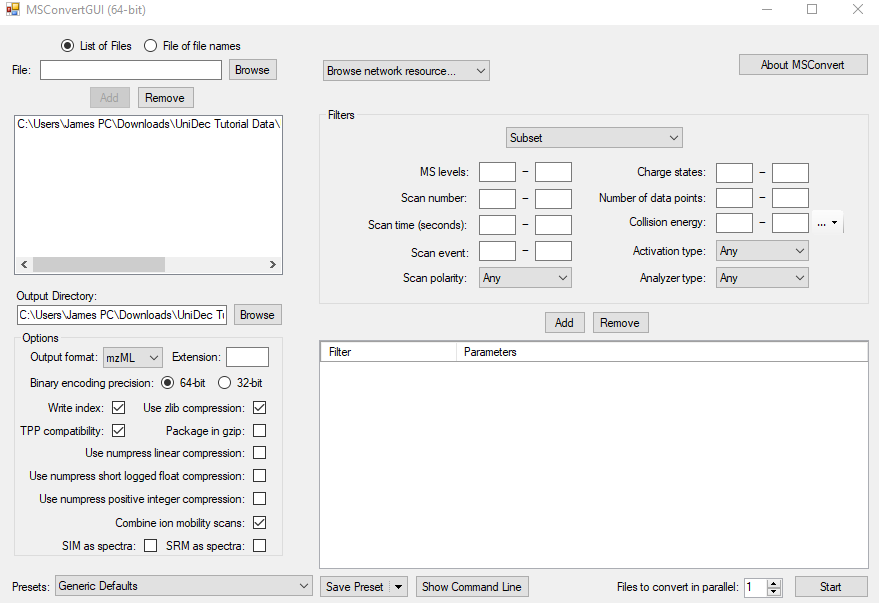
Next, we need to select the output format, and the selections should be as follows: mzML format, 64-bit encoding, write index, TPP compatibility, use zlib compression, and most importantly **COMBINE ION MOBILITY SCANS.** If you do not select this last feature, the file will no longer contain any IMS information. For non-IMS Agilent users, this last feature should be omitted from your output selections. Finally, under the “Filters” section, leave the drop down selection as “Subset”, input the determined scan times (for this tutorial 530-575), and hit the “add” button.

Finally, hit the “start” button in the lower right corner of the window, which will cause the following pop-up showing the progress of the conversion:

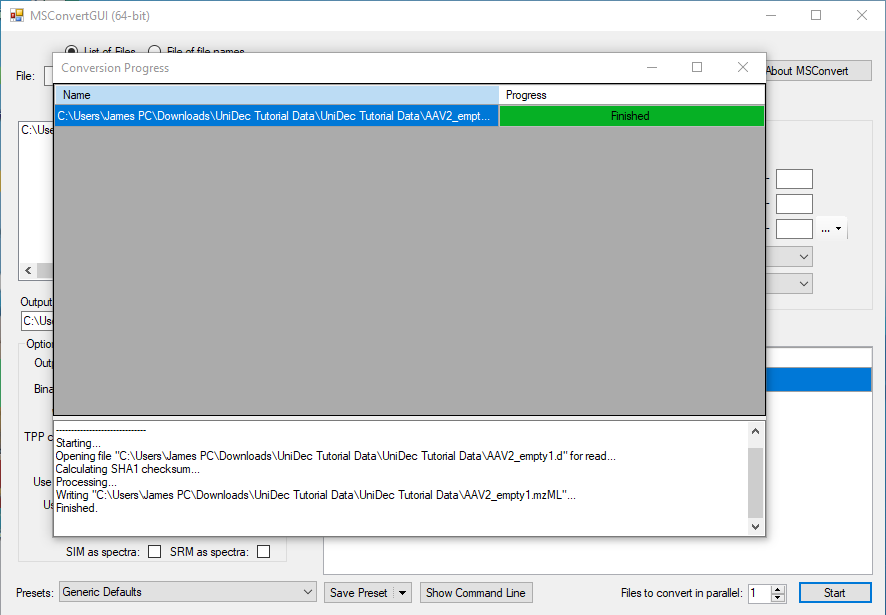

### Downloading and Running UniDec

Following successful file conversion, download the UniDec ZIP files at^2^: [**https://github.com/michaelmarty/UniDec/releases**](https://github.com/michaelmarty/UniDec/releases)

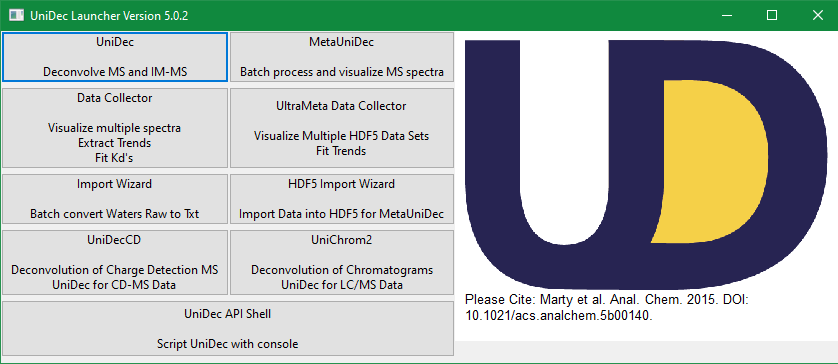
Once the folder has downloaded, extract it, open it, and scroll to the file labelled “GUI_UniDec.” Double click on this icon to launch the UniDec app. *Note – it may be necessary to right click on the file and select “run as administrator” depending upon the safety settings present on your computer.

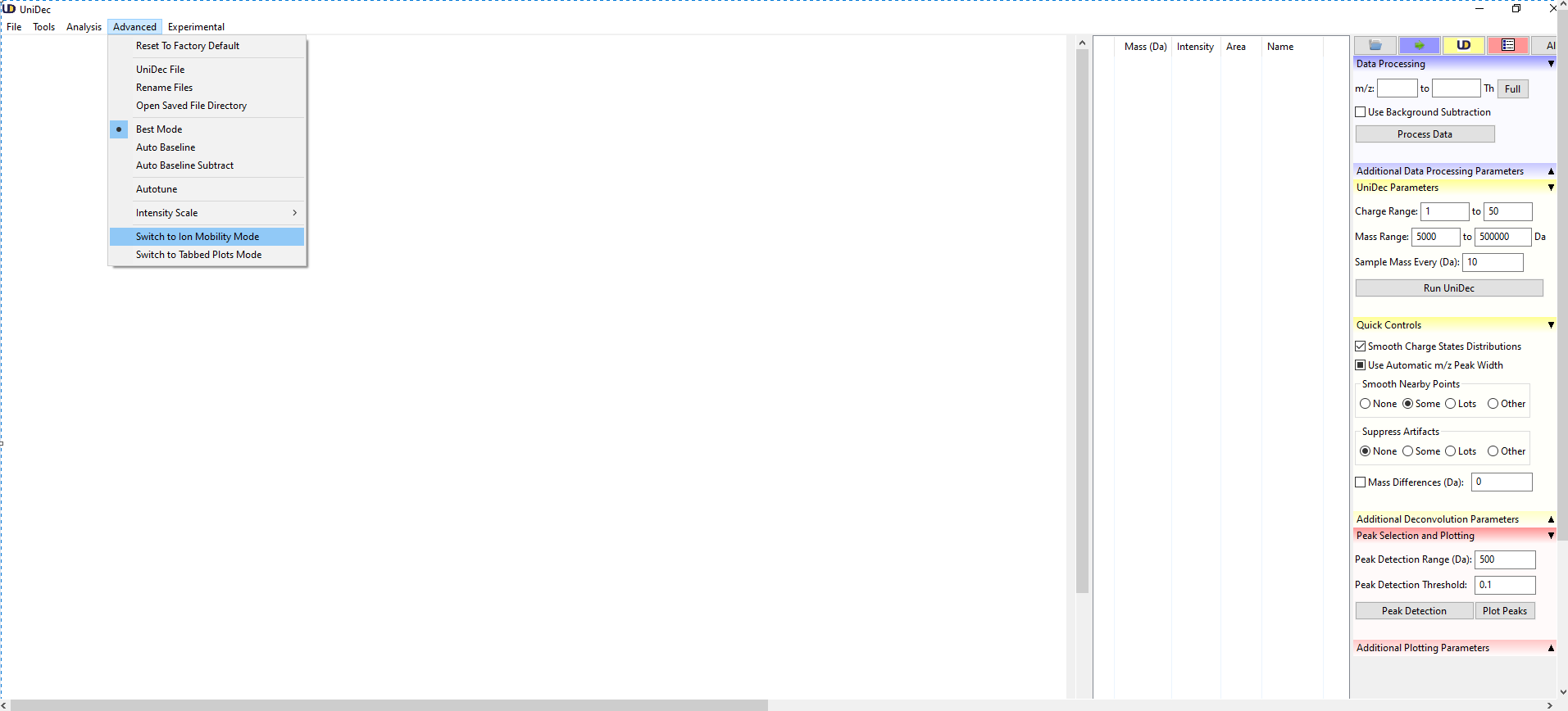
Begin by clicking on the top left button “UniDec – Deconvolve MS and IM-MS” which will open the program. For IMS data, go to the “Advanced” tab at the top left of the program, and select “switch to ion mobility mode” in the dropdown menu. Note: it may take UniDec a few minutes to initialize and run, so do not be discouraged if a “not responding” message pops up, simply click into the UniDec window and give the program a few minutes to catch up!

After switching into ion mobility mode, go to the “file” tab and select “Open File (Text, mzML, or Thermo RAW).” This will open another window, where you should go to the folder you made containing the mzML versions of your data files, and select the desired file.

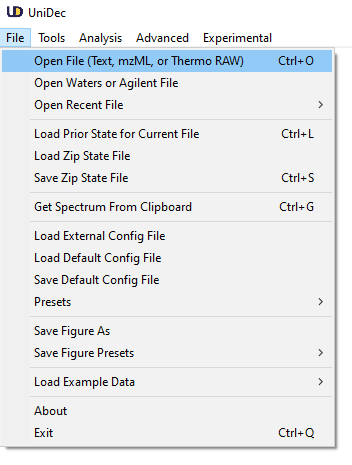

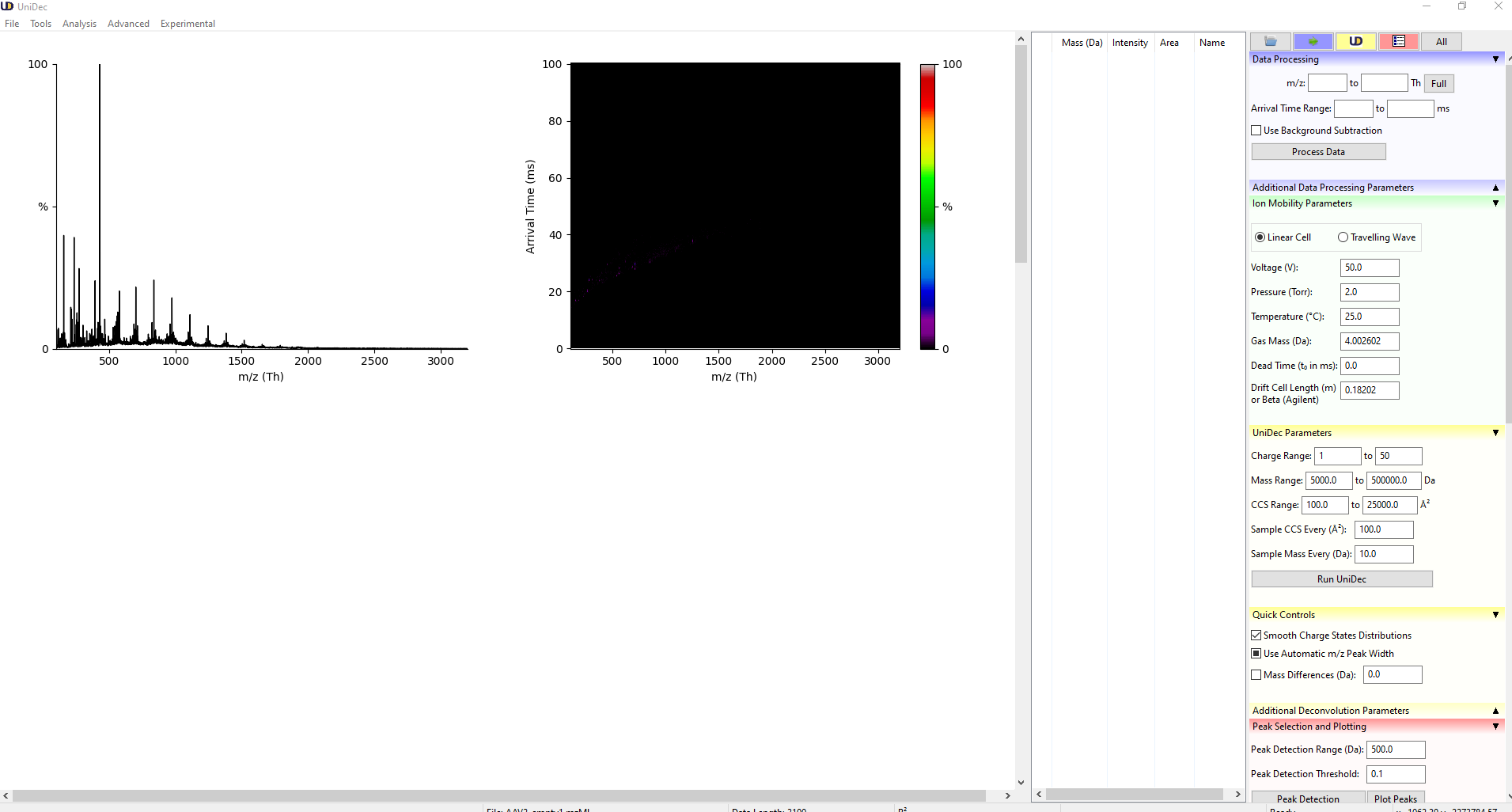
After opening the file, it will create the following plots in UniDec:

Following the successful opening of your file, you are free to alter your data processing, ion mobility, and UniDec parameters to suit your analysis needs.

For these analyses, minimal user interference was used, with data processing parameters being adjusted only to remove empty space in the *m/z* and arrival time dimensions, as shown below:

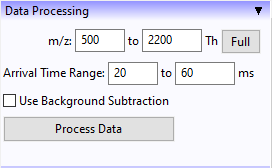

An important note is that for Agilent IMS collected using the single-field method, it is necessary to make the following alterations to your parameters. Linear cell should be selected, and voltage, pressure, and temperature should all be set to 0. Gas mass should be representative of your drift cell buffer gas, in this case nitrogen gives a mass of 28.006 Da. The last two parameters, dead time (t_o_ for us) and drift cell length (β for us) come from the calibration curve generated by measuring the ions in the Agilent tune mix. For these experiments, the values obtained from the tune calibration curve are t_o_ = -0.37206756, and β =
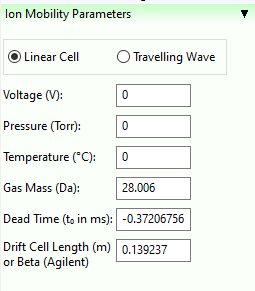
0.139237.

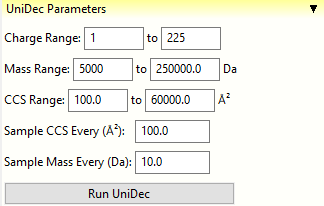
Depending upon the specific analyte you are looking to analyze, you can change your UniDec parameters to the appropriate charge range, mass range, and CCS range. For these analyses, the charge range was set to 1+ to 225+, the mass range was 5,000 Da to 250,000 Da, and the CCS range was 100 Å^2^ to 60,000 Å^2^.

After all parameters have been optimized for your data, hit “Run UniDec” to start the process. Following successful deconvolution, UniDec will create a variety of plots including: mass spectrum showing raw data vs. fit data, a zero-charge mass spectrum, a charge state assigned nested spectra of arrival time vs *m/z*, assigned charge state vs. *m/z*, CCS vs mass, and CCS vs. charge. To create UniDec’s 4-dimensional cube plots, scroll to the bottom of the right hand side of the screen, and click the “additional plotting parameters” drop down. Feel free to choose your own 2D and peak color maps, as well as your own normalization technique if desired. It is recommended to keep “discrete plot” and “publication mode” selected, as well as “reconvolved/profile.” The “marker threshold” and “species separation” were kept at the default values of 0.1 and 0.025 respectively. An example of produced 4-D plots can be seen below:

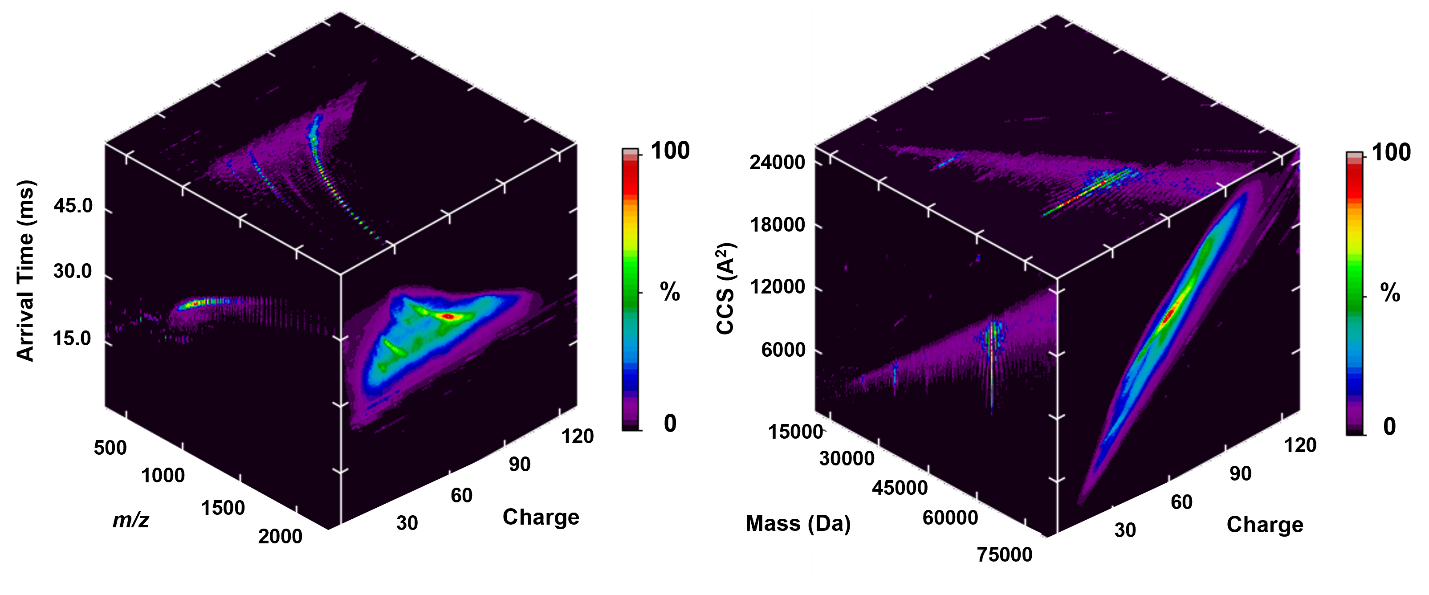

### Conclusion

In this tutorial, you have learned how to convert Agilent .d data files to .mzML files using MSConvert, a necessary conversion to facilitate analysis and visualization using UniDec. You now understand the settings required for MSConvert to include IMS information in the produced .mzML file. You have learned to run and optimize UniDec for IMS-MS data, including the production of 4-dimensional cube plots. With the understanding of how to convert and upload .d files to UniDec with MSConvert, you can now begin to utilize UniDec as a tool to help deconvolute, visualize, and analyze complex MS and IMS-MS data files collected on Agilent platforms.
