## Supplementary material for "Characterizing Adeno-Associated Virus Capsids with both Denaturing and Intact Analysis Methods": AAV Supplementary Methods

***Disclaimer***

Certain commercial equipment, software, or materials are identified in this paper to specify the experimental procedure adequately. Such identification is not intended to imply recommendation or endorsement by the National Institute of Standards and Technology, nor is it intended to imply that the equipment, software, or materials identified are necessarily the best available for the purpose.

***Comments on IM-MS Data presented in this Work***

In this supporting information we detail instrument settings used to collect LC-IMS-MS data in this manuscript, and provide raw supporting spectra for AAVs and VP subunits as noted below.

**Table of Contents**

**S-3** – Figure S1 Supplemental total ion chromatograms of empty and denatured AAV2, AAV6, and AAV8

**S-4** – Figure S2: Comparison of the total ion chromatograms of empty and filled AAVs.

**S-5** – Figure S3: Overlay of drift spectra for AAV2, AAV6, and AAV8 at varying charge states

**S-6** – Table S1: Chromatographic conditions and gradient used in this analysis.

**S-7** – Table S2: Source conditions used in this analysis.

**S-8** – Table S3: IM-QTOF parameters.

**S-9** – Table S4: UniDec parameters for the deconvolution and visualization of LC-IMS-MS data

**S-10** – Table S5: CCS calculations for unmodified and post-translationally modified VP1 for AAV2, AAV6, and AAV8.

**S-11** – Table S6: CCS calculations for unmodified and post-translationally modified VP3 for AAV2, AAV6, and AAV8.

**S-12** – Table S7: CCS calculations for unmodified and post-translationally modified VP2 for AAV2, AAV6, and AAV8.

**S-13** – Table S8: ΔCCS (%) for comparisons of VP3 for AAV2, AAV6, and AA8.

**S-14** – Table S9: ΔCCS (%) for comparisons of VP1 for AAV2, AAV6, and AA8.

**S-15** – Table S10: ΔCCS (%) for comparisons of VP2 for AAV2, AAV6, and AA8.

**S-16** – Figure S4: UniDec CD plots of for empty AAV2, AAV6, and AAV8

**S-17** – Figure S5: UniDec CD plots of for filled AAV2, AAV6, and AAV8

**S-18** – Figure S6: UniDec CD plots of for mixed AAV2, AAV6, and AAV8

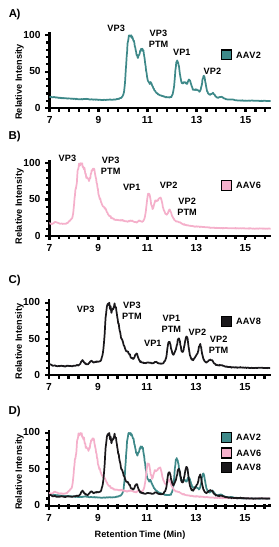

**Supplemental Figure S1.** Total ion chromatograms (TICs) of empty and denatured A) AAV2, B) AAV6, C) AAV8, and D) overlay of all three. TIC of denatured AAV2 (*A*) shows separation of all three VP subunits, as well as post-translationally modified (PTM) VP3 (oxidation). TIC of AAV6 (*B)* shows similar separation of VP subunits, as well as separation of VPs 3 and 2 with their modified peaks (oxidation). TIC of AAV8 (*C*) shows separation of all VP subunits from each other, as well as from their PTM peaks, which were observed in all 3 subunits as a phosphorylation. Panel (*D)* shows an overlay of the TICs for AAV2, AAV6, and AAV8.

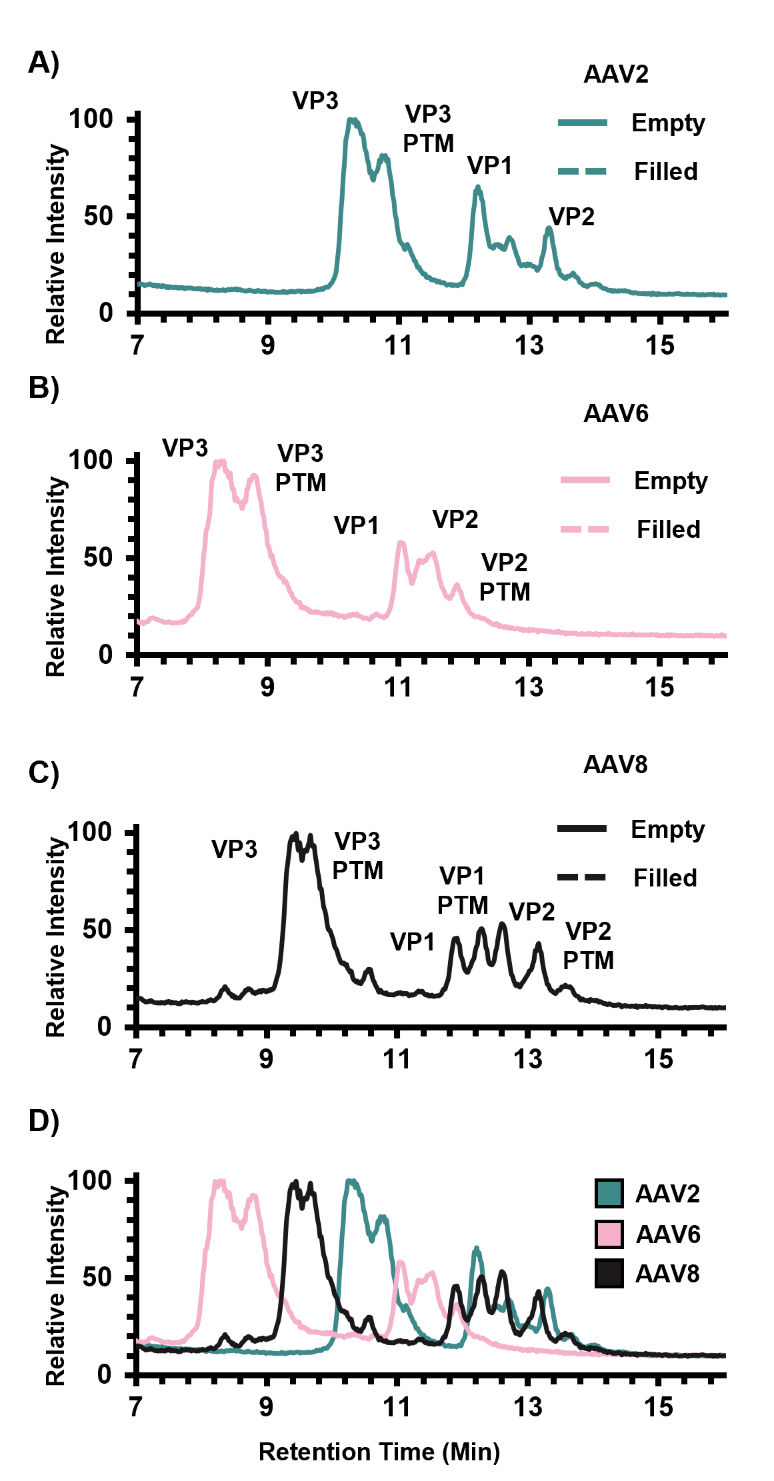

**Supplemental Figure S2.** Total ion chromatogram (TICs) overlays of empty and filled denatured A) AAV2, B) AAV6, C) AAV8. TIC of denatured AAV2 (*A*) shows separation of all three VP subunits, as well as post-translationally modified (PTM) VP3 (oxidation). TIC of AAV6 (*B)* shows similar separation of VP subunits, as well as separation of VPs 3 and 2 with their modified peaks (oxidation). TIC of AAV8 (*C*) shows separation of all VP subunits from each other, as well as from their PTM peaks, which were observed in all 3 subunits as a phosphorylation.

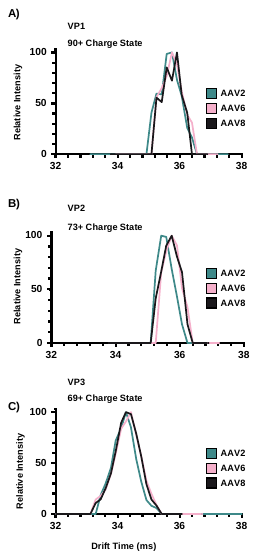

**Supplemental Figure S3.** Overlay of drift spectra for AAV2, AAV6, and AAV8. Three different charge states representing the low (90+), mid (73+) and high (69+) *m/z* values observed in the experimental spectra. Jagged AAV8 peak in the 90+ comparison arose from low abundance of the AAV8 90+ charge state. Overlap in drift spectra illustrates consistency of CCS values across the three AAV serotypes.
**Supplemental Table S1.** Chromatographic conditions and gradient used in this analysis. Method adapted from Liu, A. *et al. J. Pharmaceutical and Biomedical Anal.* **2020.** *189,* 113481.

**A)**

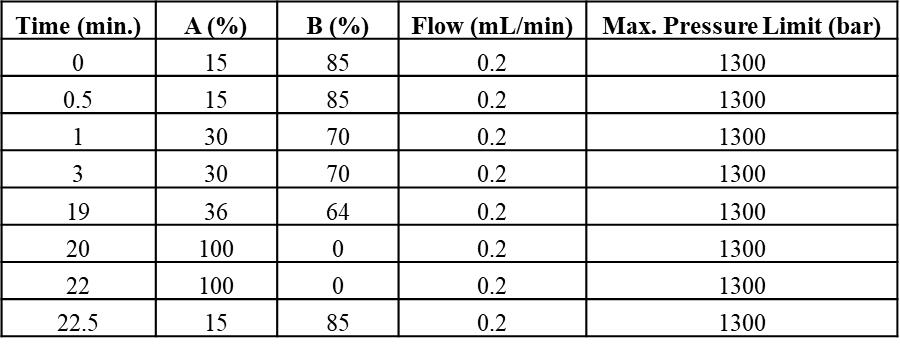

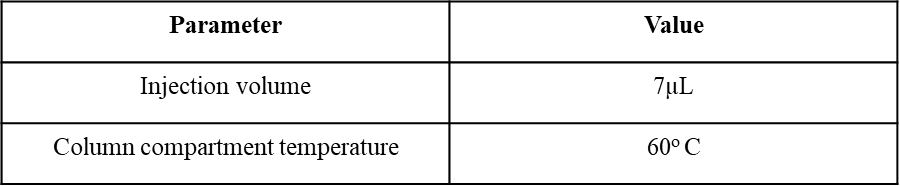
**Supplemental Table S2.** Positive ionization mode electrospray ionization source parameters used for this analysis. For conversion, 1 psi = 6894.76 Pa

**B)**

| **Parameter** | **Value** |
| --- | --- |
| Ionization Mode | Positive |
| Gas Temperature | 350^o^ C |
| Drying Gas | 8 L/min |
| Nebulizer | 15 psi |
| Sheath Gas Temperature | 275^o^ C |
| Sheath Gas Flow | 11 L/min |
| Vcap | 4000 V |
| Nozzle Voltage | 2000 V |
| Fragmentor | 400 V |
| Oct 1 RF Vpp | 750 V |

**Supplemental Table S3.** IM-QTOF MS parameters used.

| **Parameter** | **Value** |
| --- | --- |
| Min. Mass Range | 50 *m/z* |
| Max. Mass Range | 3200 *m/z* |
| Frame Rate | 0.9 Frames/s |
| IM Transient Rate | 10 IM Transients/Frame |
| Max. Drift Time | 60 ms |
| TOF Transient Rate | 600 Transients/IM Transient |
| IM Trap Fill Time | 40000 µs |
| IM Trap Release Time | 200 µs |

**Supplemental Table S4:** UniDec parameters used for the denatured AAVs, including data. processing (A), ion mobility (B), and UniDec (C) parameters. Beta and t_o_ are from our single-field calibration sheet and are specific to this experiment. Values come from calibration curve generated from the drift times and CCS values of the Agilent tune mix. Please note that the temperatures, pressures, and voltages highlighted in part B are not reflective of the actual ion mobility parameters used in the data acquisition; however, Unidec code dictates that they should be given a value of zero as the beta and Tfix terms provide the CCS calibration from analyte drift times. For conversion, 1 Torr = 133.322 Pa and 1Da = 1.660E-27 kg.

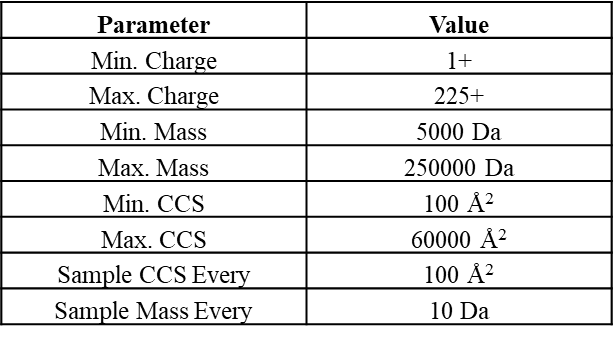

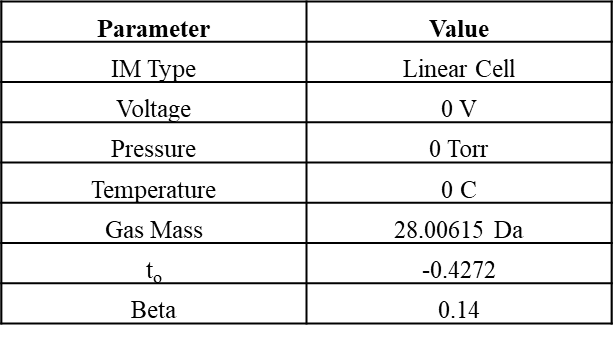

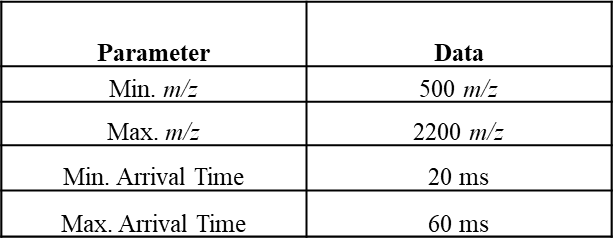
**Supplemental Table S5**. Average CCS values calculated for each serotype at the specified charge states for A) unmodified VP1, and B) Post-translationally modified VP1 in AAV2, AAV6, and AAV8.

**C)**

**UniDec Parameters**

**B)**

**Ion Mobility Parameters**

**A)**

**Data Processing Parameters**

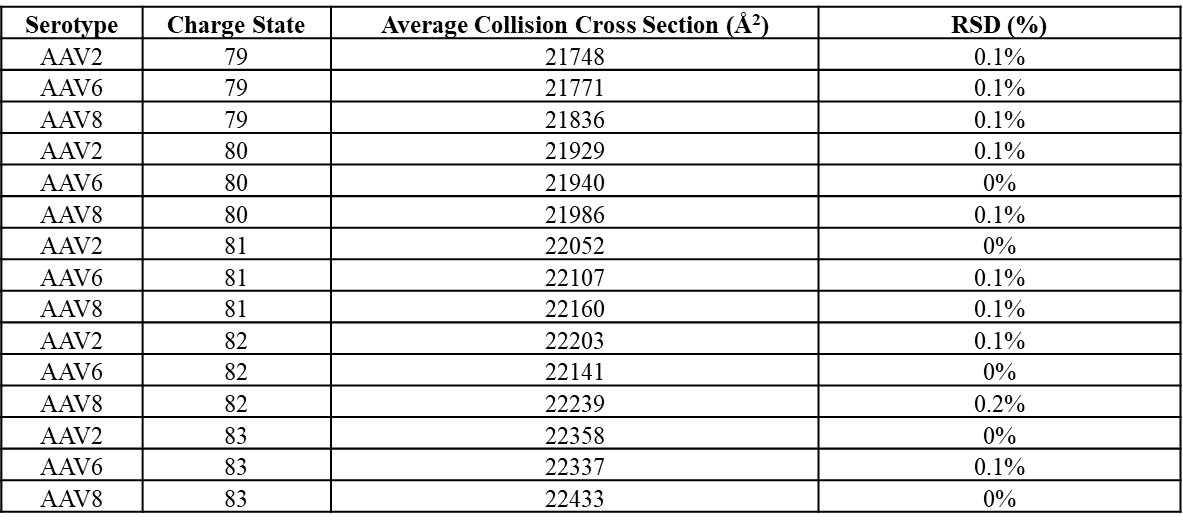

**A)**

**Unmodified VP1 Calculations**

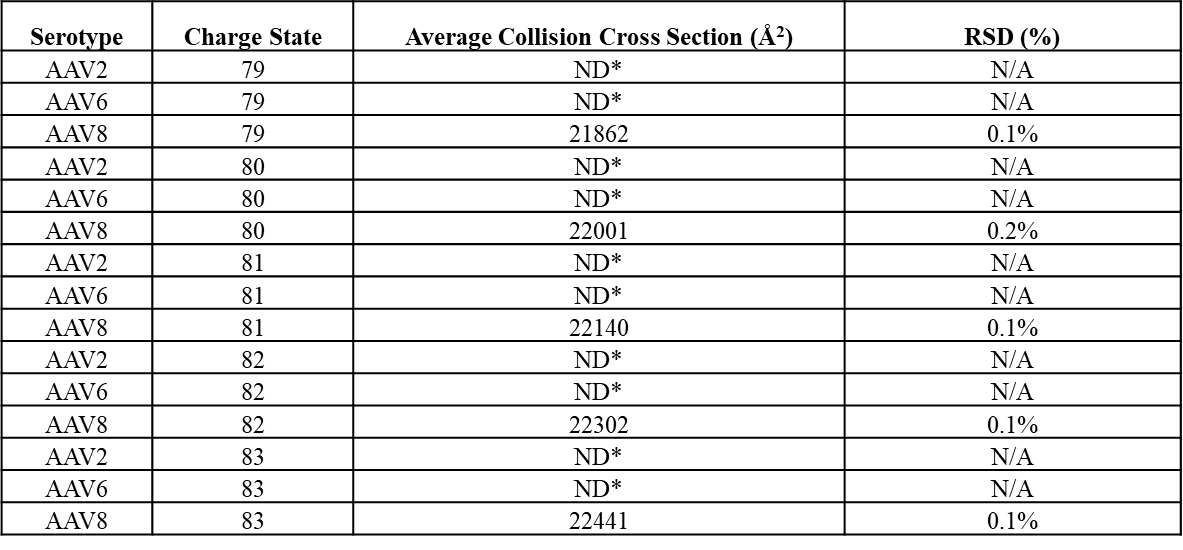

**B)**

**Post-translationally Modified VP1 Calculations**

*ND = not detected, N/A = corresponds to inability to calculate RSD (%) due to non-detection of analyte

**Supplemental Table S6:** Average CCS values calculated for each serotype at the specified charge states for A) unmodified VP3, and B) Post-translationally modified VP3 in AAV2, AAV6, and AAV8.

**A)**

**Unmodified VP3 Calculations**

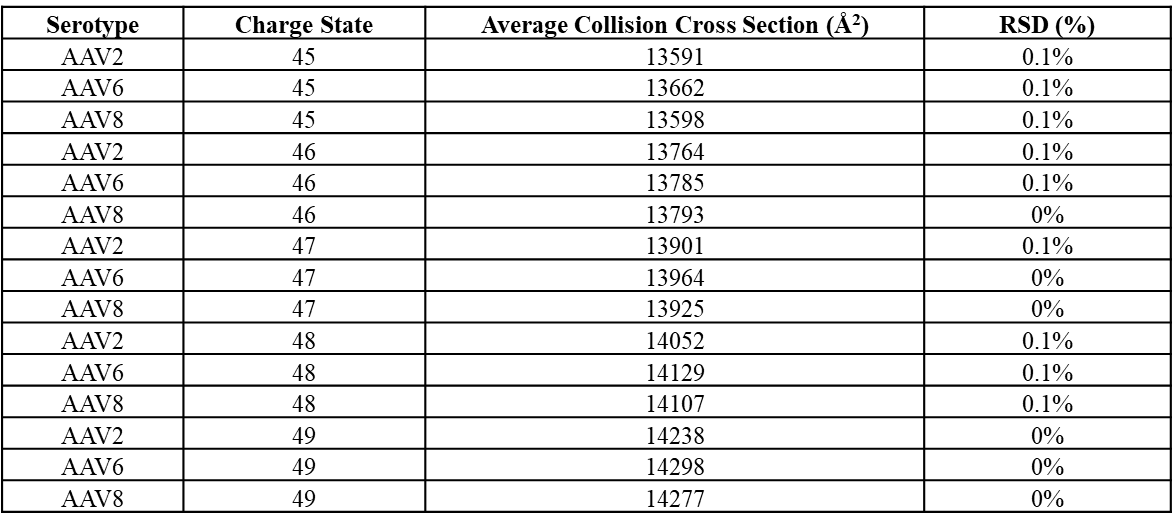

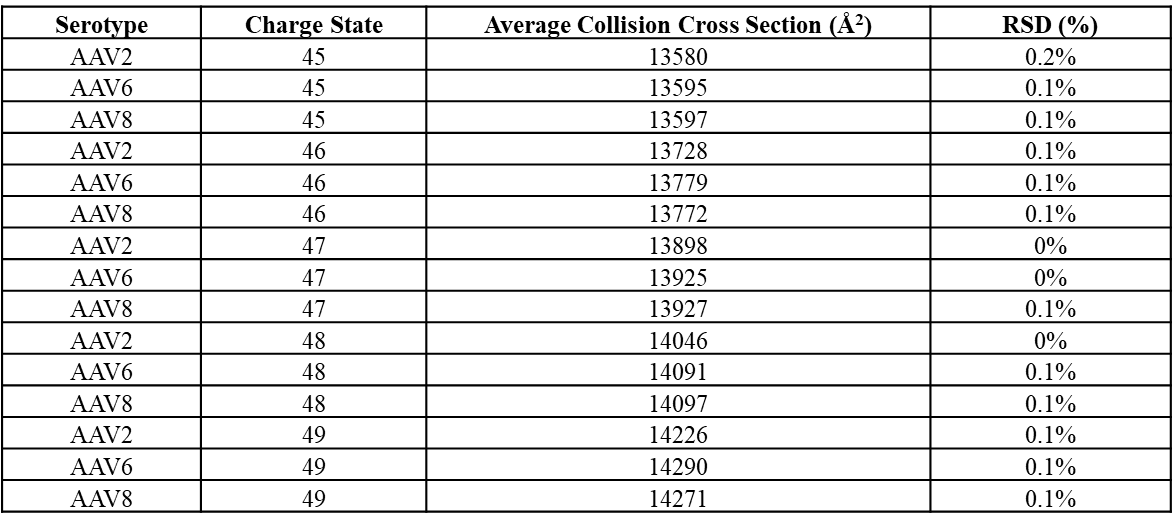

**B)**

**Post-translationally Modified VP3 Calculations**

**Supplemental Table S7**. Average CCS values calculated for each serotype at the specified charge states for A) unmodified VP2, and B) Post-translationally modified VP2 in AAV2, AAV6, and AAV8.

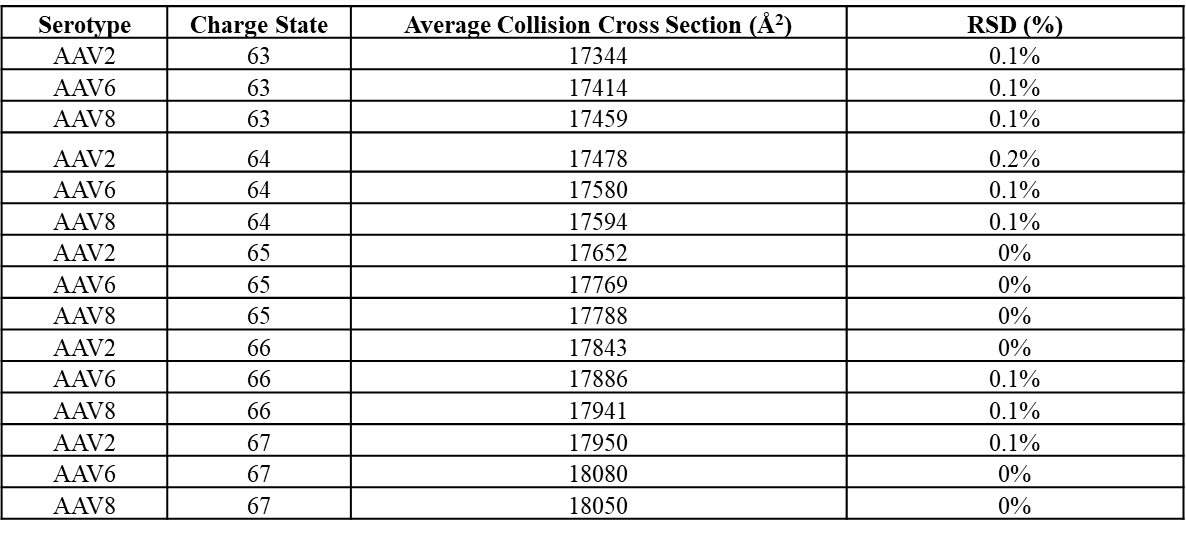

**A)**

**Unmodified VP2 Calculations**

**B)**

**Post-translationally Modified VP2 Calculations**

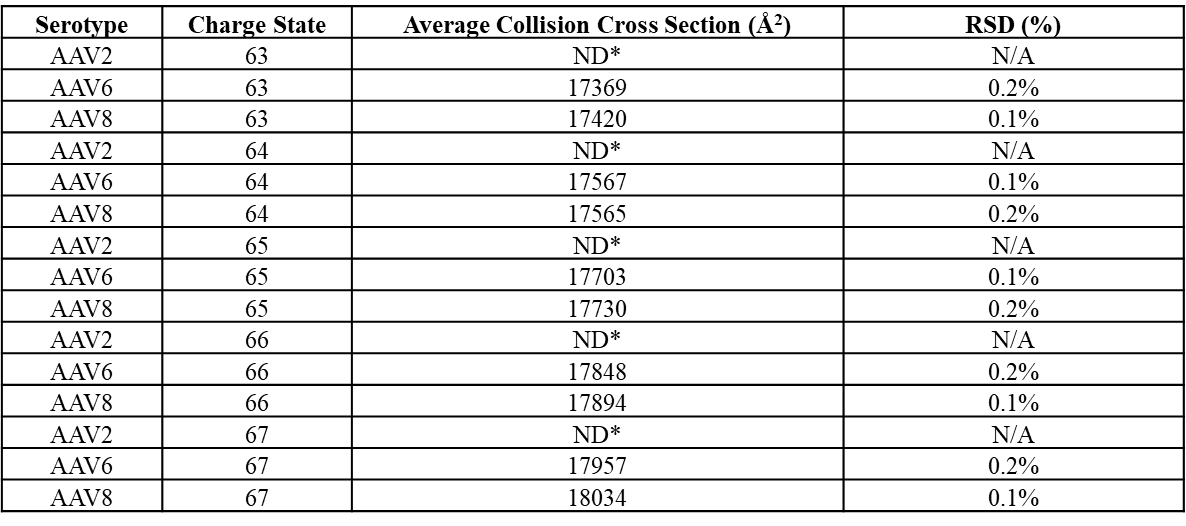

*ND = not detected, N/A = corresponds to inability to calculate RSD (%) due to non-detection of analyte

**Supplemental Table S8**. ΔCCS calculations for given comparison at specified charge states. For our platform (Agilent 6560, *R*_p_ ~60 for singly charged ions) a ΔCCS (%) of ~2.5-3% is needed for ~50% separation.

**VP3 Collision Cross Section Comparisons**

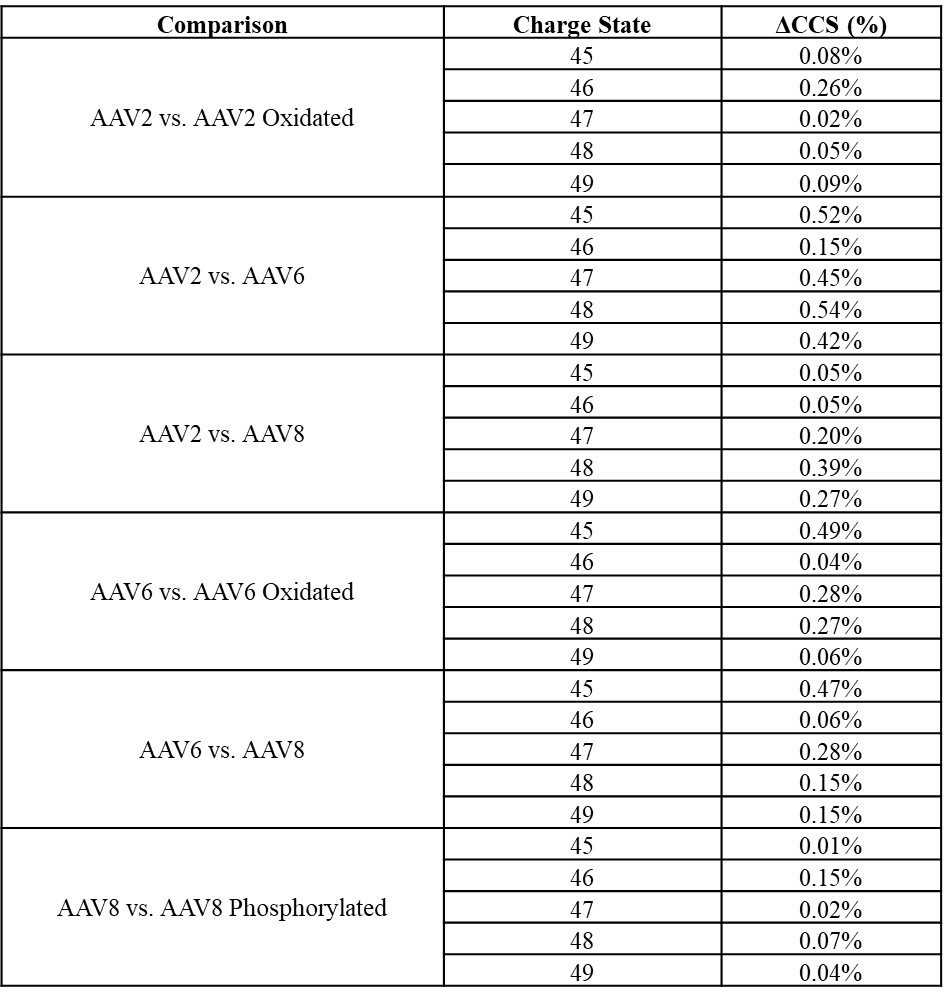

**Supplemental Table S9**. ΔCCS calculations for given comparison at specified charge states. For our platform (Agilent 6560, *R*_p_ ~60 for singly charged ions) a ΔCCS (%) of ~2.5% - 3% is needed for ~50% separation.

**VP1 Collision Cross Section Comparisons**

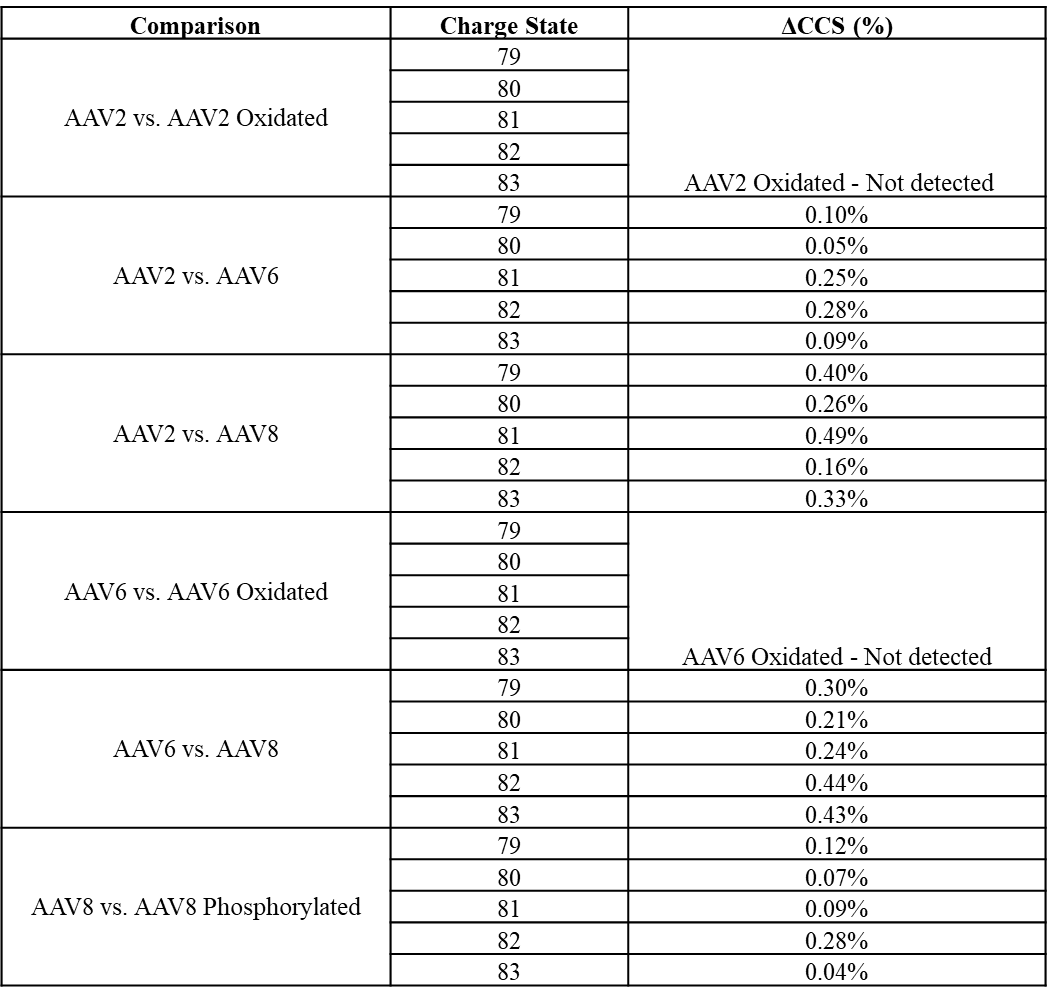

**Supplemental Table S10**. ΔCCS calculations for given comparison at specified charge states. For our platform (Agilent 6560, *R*_p_ ~60 for singly charged ions) a ΔCCS (%) of ~2.5-3% is needed for ~50% separation.

**VP2 Collision Cross Section Comparisons**

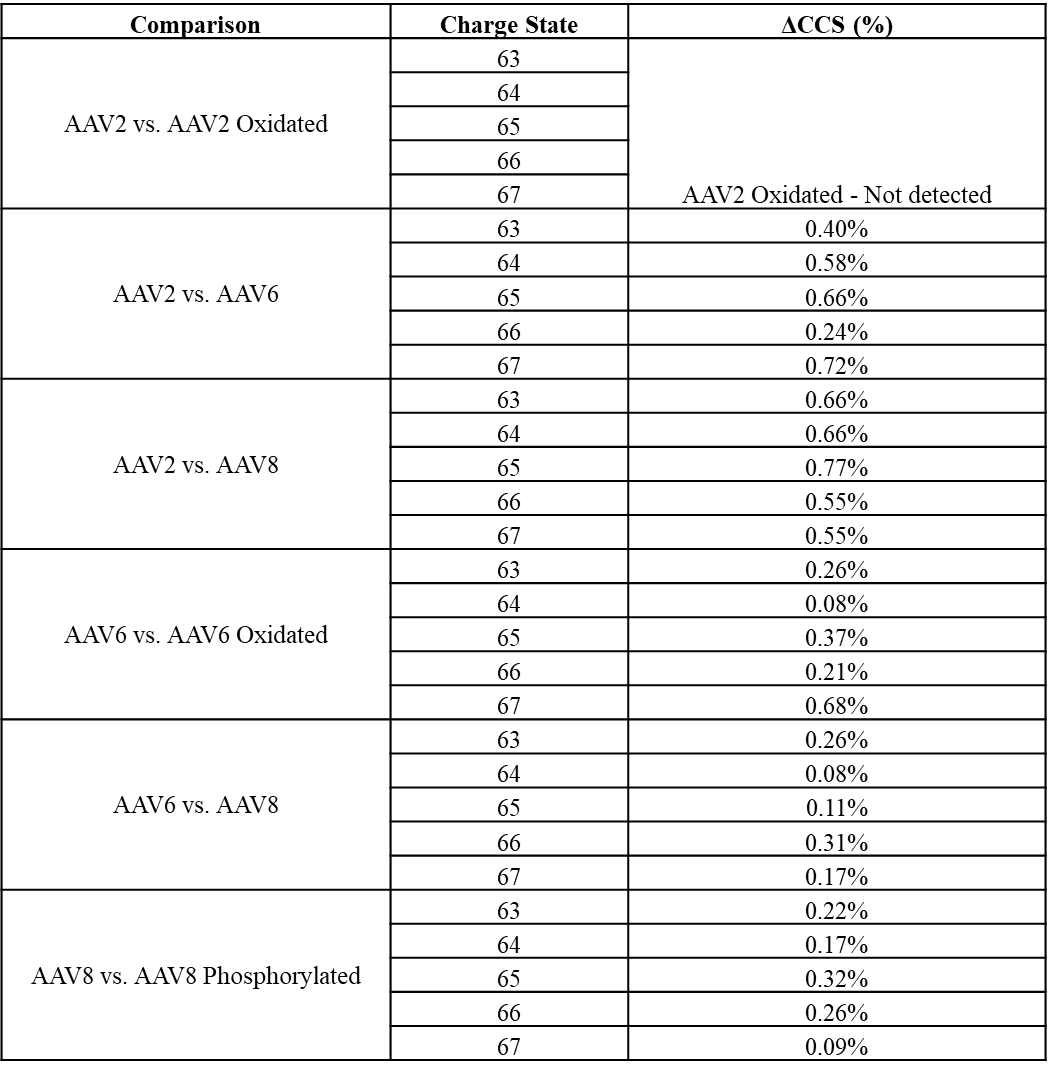

**Supplemental Figure S4**. UniDecCD plots showing charge vs *m/z* and charge vs mass (MDa) for empty AAV capsids, for AAV2, AAV6, and AAV8.

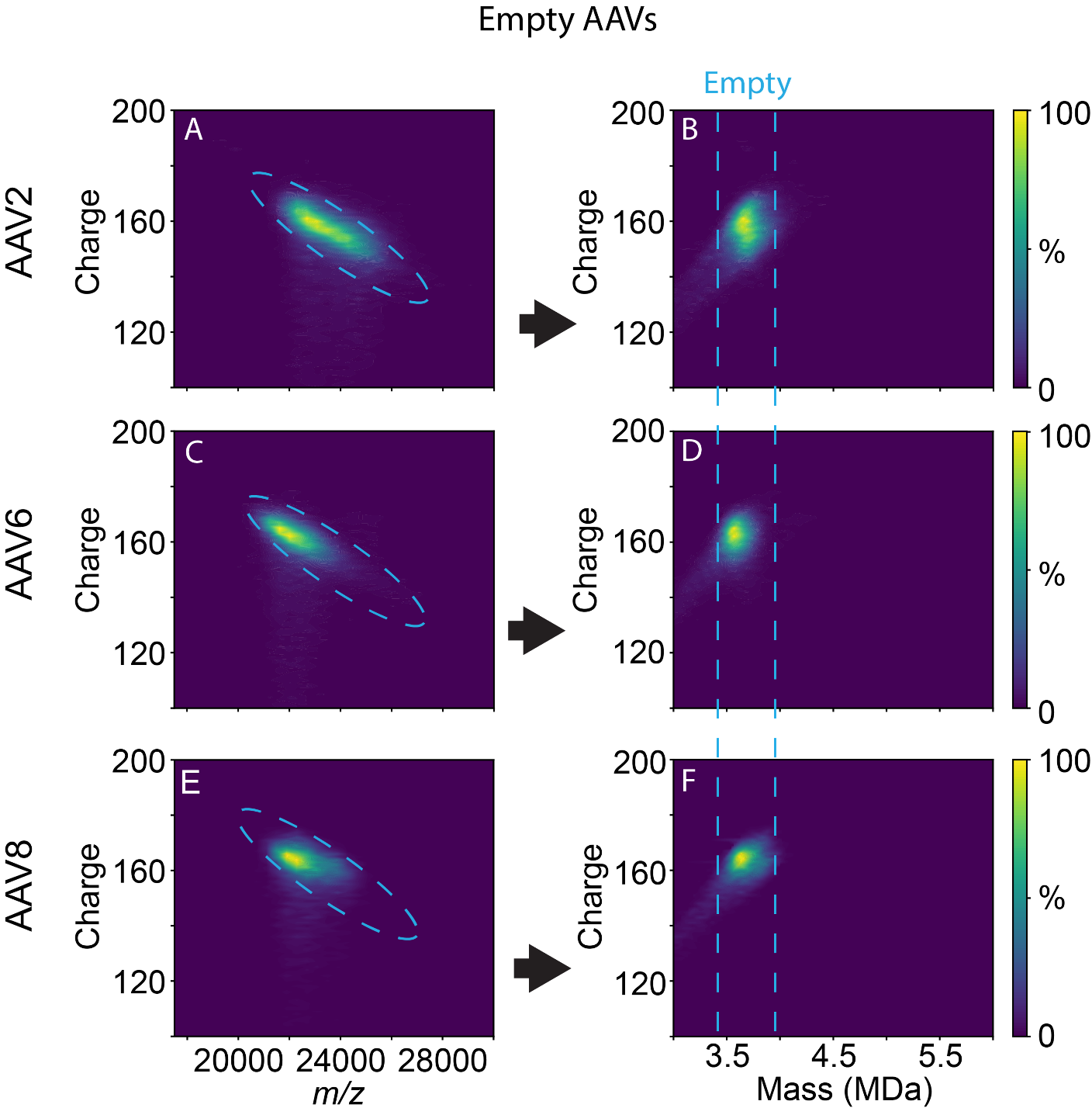

**Supplemental Figure S5**. UniDecCD plots showing charge vs *m/z* and charge vs mass (MDa) for filled AAV capsids, for AAV2, AAV6, and AAV8.

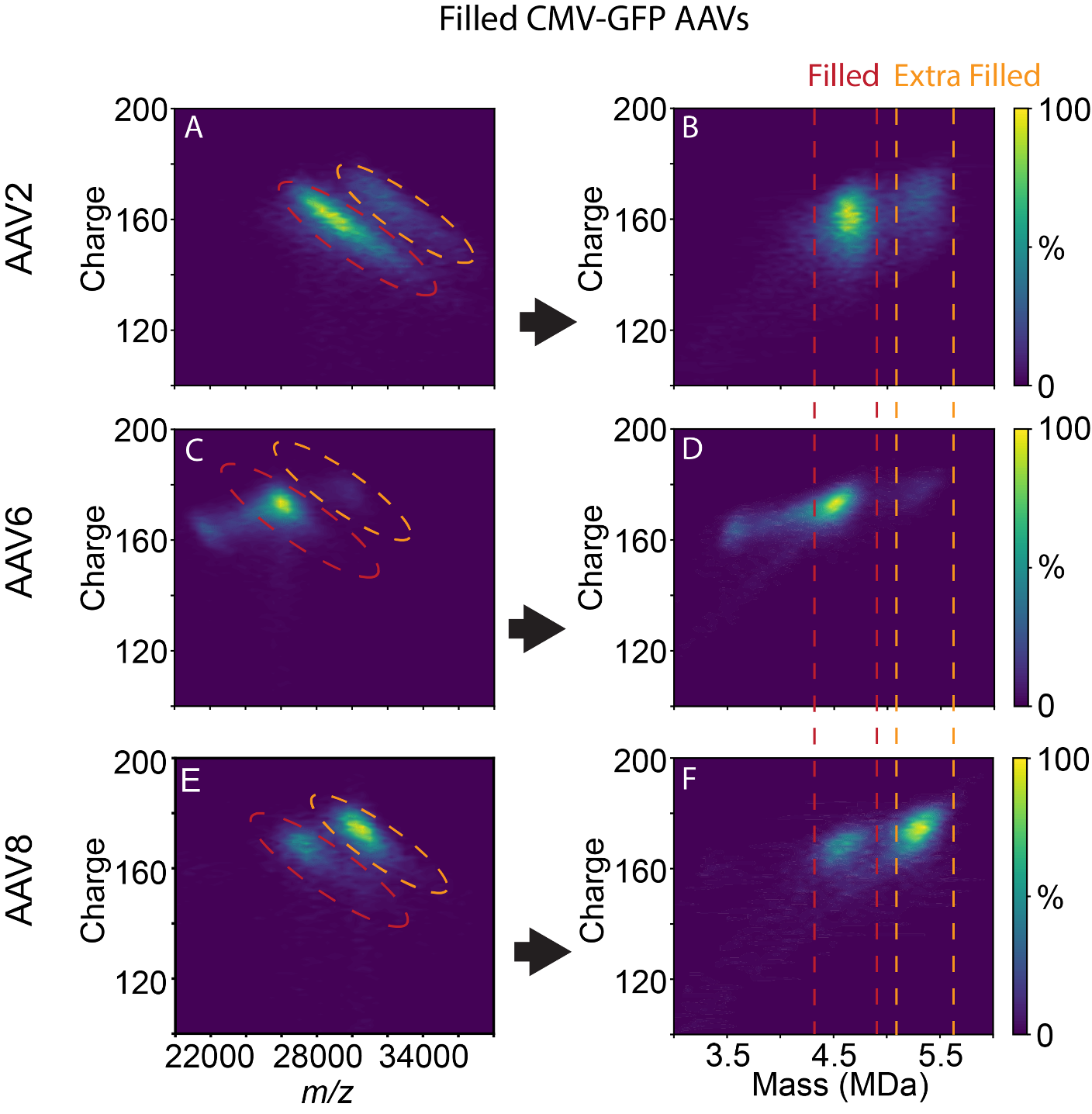

**Supplemental Figure S6**. UniDecCD plots showing charge vs *m/z* and charge vs mass (MDa) for mixed AAV capsids, for AAV2, AAV6, and AAV8.

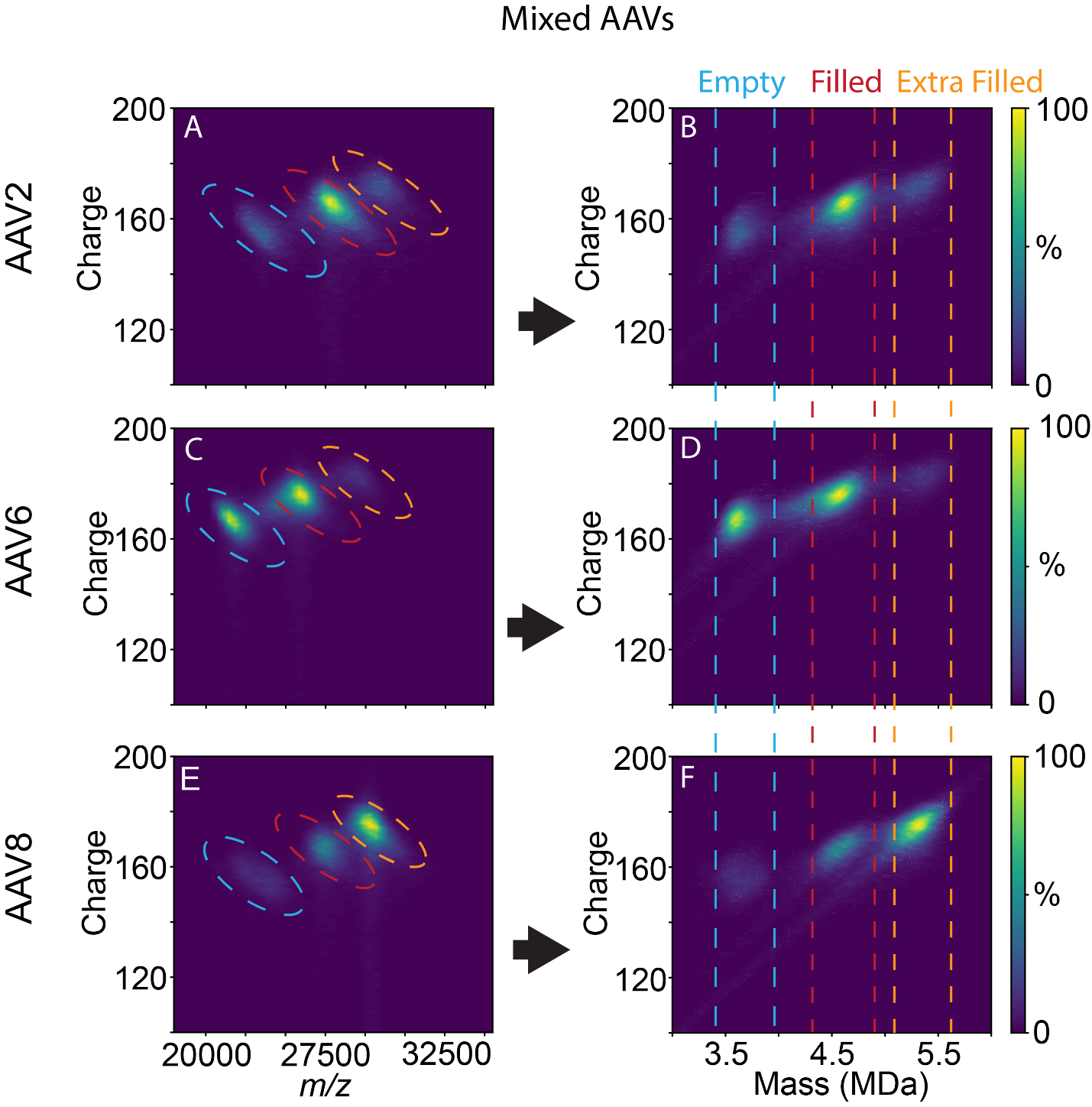
